# Potentiating CD8^+^ T cell antitumor activity by targeting the PCSK9/LDLR axis

**DOI:** 10.1101/2020.12.03.403121

**Authors:** Juanjuan Yuan, Ting Cai, Xiaojun Zheng, Yangzi Ren, Jinwen Qi, Xiaofei Lu, Huihui Chen, Huizhen Lin, Zijie Chen, Mengnan Liu, Shangwen He, Qijun Chen, Siyang Feng, Yinjun Wu, Zhenhai Zhang, Yanqing Ding, Wei Yang

## Abstract

Metabolic regulation has proven to play a critical role in T cell antitumor immunity. Cholesterol metabolism is a key component of this response but remains largely unexplored. Herein, we found that the LDL receptor (LDLR), which has been previously identified as a transporter for cholesterol and fatty acids, plays a pivotal role in regulating CD8^+^ T cell antitumor activity, with the genetic ablation of LDLR significantly attenuating CD8^+^ T cell activation and clonal expansion. Additionally, we found that LDLR interacts with the T-cell receptor (TCR) signalosome and regulates TCR signaling, facilitating CD8^+^ T cell activation and effector function. Furthermore, we found that the tumor microenvironment downregulates CD8^+^ T cell LDLR levels and TCR signaling via tumor cell-derived PCSK9, which binds and prevents the recycling of LDLR and TCR into the plasma membrane. Our findings indicate that genetic deletion or pharmacological inhibition of PCSK9 in tumor cells can enhance the antitumor activity of CD8^+^ T cells by alleviating the tumor microenvironment’s suppressive effect on CD8^+^ T cells and consequently inhibit tumor progression. While previously established as a hyperlipidemia target, this study highlights PCSK9 as a potential target for cancer immunotherapy as well.

## INTRODUCTION

CD8^+^ T cell-based immunotherapy ha s emerged as one of the most important cancer therapeutic strategies. Particular success has been seen with new approaches like immune checkpoint blockade (ICB), targeting PD-1 and CTLA4, and chimeric antigen receptor T (CAR-T) cell therapy, both of which have been approved for the treatment of a variety of cancers (Leach et al., 1996; Morgan et al., 2006; Wolchok et al., 2013; Maude et al., 2014). Despite the clinical successes, the efficacy of CD8^+^T cell-based immunotherapy varies substantially across malignancies and individuals (Maus et al., 2013; Rizvi et al., 2015; Neelapu et al., 2018; Rafiq et al., 2020). As such, further investigation into the regulatory mechanisms and efficacy factors of immune therapy is warranted.

Mechanistically, once activated by tumor antigens peripheralCD8^+^ T cells will traffic to the tumor microenvironment (TME) and mediate antitumor responses (Borst et al., 2018). However, the TME possesses numerous immunosuppressive properties, primarily mediated by immune suppressive stromal cells, myeloid cells, lymphoid cells, and tumor cells themselves, limiting the antitumor activity of CD8^+^ T cells. While these immunosuppressive cells are the main cause of immunotherapy failure (Draghiciu et al., 2015; Kalluri, 2016; Kumar et al., 2017; Mantovani et al., 2017; Togashi et al., 2019), a lack of nutrients—such as glucose and lipids—as well as hypoxia in the TME are also correlated with CD8^+^ T cell dysfunction (Chang et al., 2015; Bunse et al., 2018; Gourdin et al., 2018; Leone et al., 2019; Baumann et al., 2020). These previous studies suggest that the metabolic regulation by the TME plays a critical role in CD8^+^ T cell suppression (Sukumar et al., 2013; Ho et al., 2015; Patsoukis et al., 2015; Zhang et al., 2017; Wang and Zou, 2020).

Metabolic regulation has been shown to play critical roles in T cell differentiation and effector function (Almeida et al., 2016; Kishton et al., 2017; Patel and Powell, 2017). As the primary component of lipids, cholesterol metabolism in particular is essential for CD8^+^ T cell function and its reprogramming has induced significant alterations to CD8^+^ T cell activation (Kidani et al., 2013; Wang et al., 2016; Yang et al., 2016). Recent studies have also highlighted the importance of cellular cholesterol metabolism in regulating the antitumor efficacy of CD8^+^ T cells (Yang et al., 2016; Ma et al., 2018; Ma et al., 2019; Ma et al., 2020). However, the mechanisms by which the TME reprograms CD8^+^ T cell cholesterol metabolism, and to what extent this impacts tumor immune evasion, remains unknown.

To examine how cholesterol metabolism modulates T cell function in the TME, we measured the cholesterol levels of intratumoral CD8^+^ T cells. We found that intratumoral CD8^+^ T cells have reduced cholesterol levels, resulting from the LDLR (Low Density Lipoprotein Receptor) deficiency. Furthermore, we elucidated that LDLR plays a key role in T cells’ tumoricidal effects. In addition to controlling the uptake of LDL, LDLR also interacts with CD3, a subunit of the T-cell receptor (TCR) complex, modulating the TCR signaling pathway. Additionally, it has been reported previously that PCSK9 regulates the degradation of LDLR, consequently blocking cholesterol uptake (Garcia et al., 2001; Rudenko et al., 2002; Maxwell et al., 2005; Kwon et al., 2008; Poirier et al., 2008). Upon investigation, we found that PCSK9 was highly expressed in tumors and constantly released into the TME, and that this secretion of PCSK9 dampened the immune response of CD8^+^ T cells via hindering LDLR expression and ultimately inhibited TCR signaling and effector function. These findings highlight the PCSK9-LDLR regulatory network as a novel potential target in cancer immunotherapy.

## RESULTS

### LDLR deficiency hinders the antitumor activity of CD8^+^ T cells

Antigen recognition induces cholesterol metabolic reprogramming in CD8^+^ T cells, which enables the cells to acquire sufficient cholesterol to support clonal expansion and effector function (Zech et al., 2009; Kidani et al., 2013; Yang et al., 2016; Newton et al., 2018). The tumor microenvironment has been demonstrated as a hypoxia and nutrient restricted environment (Chang et al., 2015; Semenza, 2016; Zhang and Ertl, 2016; Cascone et al., 2018). Whether there is sufficient cholesterol in the TME to support CD8^+^ T cells’ antitumor activity, and if not how CD8^+^ T cells acquire sufficient cholesterol in such an environment, is unknown. To investigate, we analyzed the Apolipoprotein B (APOB) levels of clinical cancer samples and syngeneic mouse tumor samples. We found that the APOB level, which represents the LDL/cholesterol level, was significantly higher in the tumor regions than the paracancerous normal tissues (**Supplementary figure 1a-f**). In contrast, the cellular cholesterol levels of tumor infiltrating CD8^+^ T cells from MC38 tumor burdened syngeneic mice were lower than that of the splenic CD8^+^ T cells, when quantified by Filipin III staining (**Supplementary figure 1g, h**). These findings indicate that the reduced cellular cholesterol in CD8^+^ T cells may be due to an aspect of the T cells themselves.

**Figure 1.**
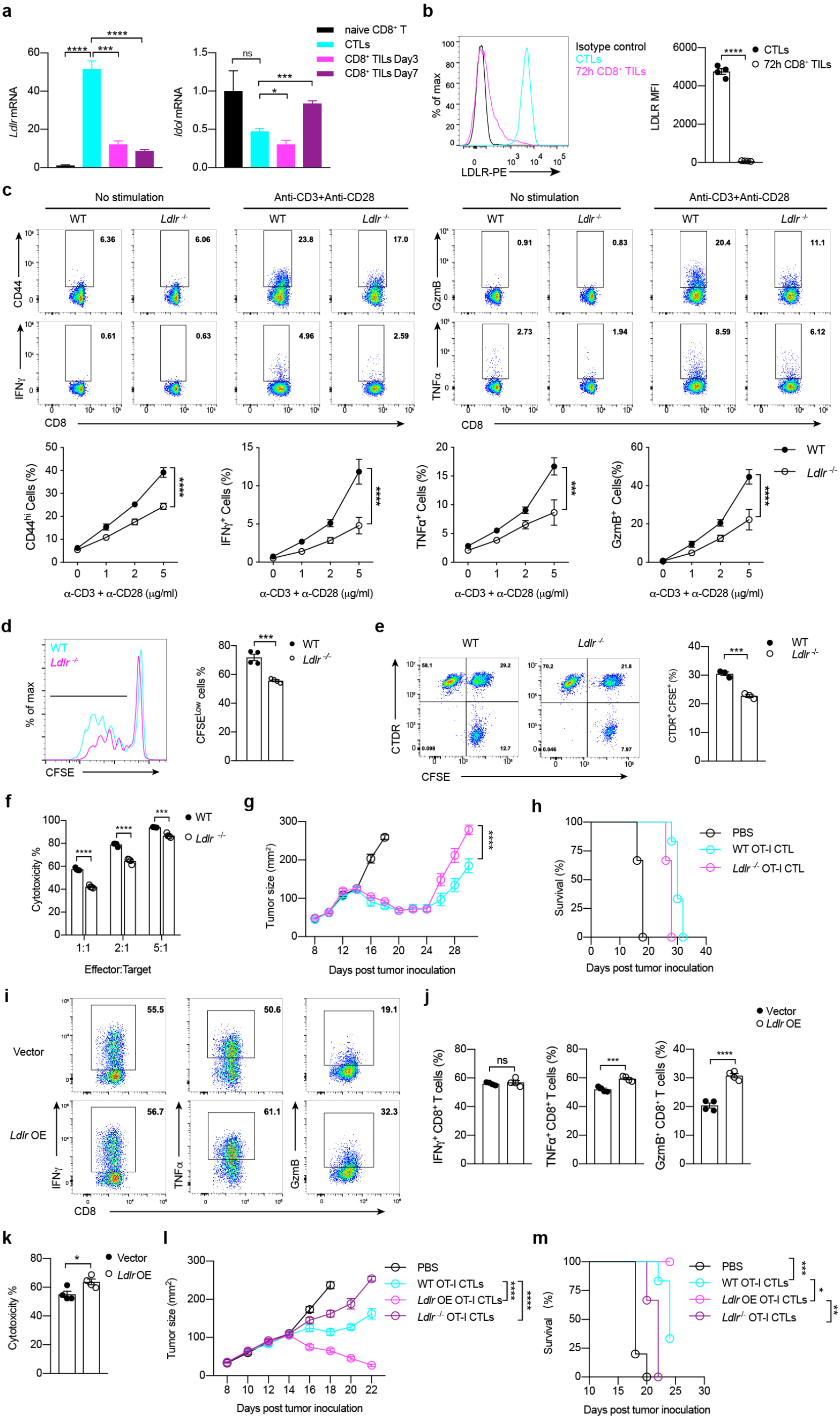
LDLR deficiency hinders the antitumor activity of CD8^+^T cells. **a**, Transcriptional level of cholesterol transport genes *Ldlr* and *Idol* in naïve CD8 T cells, CTL and CD8^+^ TILs (isolated at Day3 or Day7 post adoptive transfer), (n = 4). **b**, LDLR expression levels on CTLs and CD8^+^ TILs (isolated at 72 hours post adoptive transfer), (n = 4). **c**, T cell activation and cytokine/granule productions of WT and *Ldlr*^-/-^ CD8^+^ T cells. Naïve CD8^+^ T cells were isolated from the spleen and stimulated with anti-CD3 and anti-CD28 antibodies for 24 hours at indicated concentrations. Data were analyzed by two-way ANOVA (n = 4). **d**. CD8^+^ T cell proliferation was measured by CFSE dilution assay. CD8^+^ T cells were isolated from the spleen and stimulated with 1μg/ml plate-coated anti-CD3 and anti-CD28 antibodies for 72h (n = 4). **e**, Immunological synapse formation of WT and *Ldlr*^-/-^ CTLs. CFSE-labeled CTLs and CellTracker Deep Red (CTDR)-labeled OVA-pulsed EL4 cells were cocultured for 30 mins (n = 3). **f**, Cytotoxicity of WT and *Ldlr*^-/-^ CTLs. Splenocytes from WT and *Ldlr*^-/-^ OT-I mice were stimulated with OVA^257–264^ and IL-2 to generate mature CTLs. CTLs were incubated with OVA-pulsed CTDR-labeled EL-4 cells and CFSE-labeled non-pulsed EL-4 cells for 4 hours. The ratio of OVA-pulsed and non-pulsed EL4 cells was calculated to determine the cytotoxicity of CTLs (n = 4). **g-h**, Tumor growth (**g**) and survival (**h**) of MC38-OVA tumor-bearing *Rag2*^-/-^ mice after adoptive transfer of PBS, WT or *Ldlr*^-/-^ CTLs. Data were analyzed by two-way ANOVA (n = 6). **i-j**, Cytokine and granule productions of control and *Ldlr* OE CTLs. *Ldlr* was overexpressed in CTLs with retrovirus infection. The sorted cells were stimulated with 1μg/ml plate-coated anti-CD3 and anti-CD28 for 4 hours (n = 4). **k**, Cytotoxicity of control and *Ldlr* OE CTLs. CTLs were incubated with OVA-pulsed EL-4 cells and non-pulsed EL-4 cells for 4 hours (n = 4). **l-m**, Tumor growth (**l**) and survival (**m**) of MC38-OVA tumorbearing *Rag2*^-/-^ mice after adoptive transfer of PBS, WT, *Ldlr* OE or *Ldlr*^-/-^ CTLs. Data were analyzed by two-way ANOVA (n = 5-6). *, *P* < 0.05; **, *P* < 0.01; ***, *P* < 0.001****, *P* < 0.0001. Error bars denote for the s.e.m.

We then further evaluated the cholesterol metabolic program of tumor-infiltrating CD8^+^ T cells. In addition to reduced cholesterol biosynthesis (**Supplementary figure 1i-l**), we found that LDLR—which has been previously identified as a transporter for LDL/cholesterol (Gent and Braakman, 2004; Kwon et al., 2008)—mRNA levels in tumor infiltrating CD8^+^ T cells were much lower than in the activated cytotoxic CD8^+^ T cells (CTLs) from adoptive CTL transfer therapy in MC38 tumor burdened mice (**Figure 1a**). The reduced surface LDLR levels in tumor infiltrating CD8^+^ T cells was further validated by flow cytometric analysis (**Figure 1b**).

To determine the physiological functions of LDLR in CD8^+^ T cells, we isolated splenic CD8^+^ T cells from *Ldlr*^-/-^ mice showing normal T cell development (**Supplementary figure 1n, o**). When compared with the wild-type CD8^+^ T cells, the *Ldlr*^-/-^ CD8^+^ T cells showed impaired effector function, such as reduced cytokine and granule production, as well as lower clonal expansion rate (**Figure 1c, d**). To further assess the involvement of LDLR in the immune responses of CD8^+^ T cells *in vivo*, we generated antigen-specificand LDLR deficient CD8^+^ T cells by crossing OT-I transgenic mice and *Ldlr*^-/-^ mice.We then generated *Ldlr*^-/-^OT-I CTLs via pulsing splenocytes with OVA257-264 peptides (SIINFEKL). We found that LDLR deficiency induced the defect of immunological synapse formation (**Figure 1e**) and impaired cytotoxicity to tumor cells when we cocultured these CTLs with OVA257-264 loaded EL4 cells (**Figure 1f**). We then transferred the OT-I CTLs to the ovalbumin expressing MC38 tumor (MC38-OVA) mice. We found that *Ldlr* depletion indeed impaired the antitumor activity of CD8^+^ T cells, with the mice showing more advanced tumor progression (**Figure 1g, h**). In contrast, the overexpression of LDLR in OT-I CTLs enhanced the antitumor activity of CD8^+^ T cells both *in vitro* and *in vivo* (**Figure 1i-m**). Together, these results demonstrate that LDLR intrinsically regulates CD8^+^ T cell immune response and antitumor activity.

### The regulation of LDLR on CD8^+^ T cell effector function is not fully dependent on LDL/cholesterol

The primary function of the LDLR is to mediate the endocytosis of cholesterol enriched LDL and maintain the plasma levels of LDL (Jeon and Blacklow, 2005; Go and Mani, 2012). On the cellular level, LDL derived cholesterol is one of the resources necessary for CD8^+^ T cell proliferation and effector function (Kidani et al., 2013; Yang et al., 2016; Proto et al., 2018). To evaluate the role of LDL in CD8^+^ T cell function and further assess the function of LDLR, we first measured the LDL uptake in LDLR deficient CD8^+^ T cells. The results exhibited that LDL uptake in CD8^+^ T cells was completely dependent on LDLR (**Figure 2a**). Furthermore, we found that depleting the LDL in the medium substantially inhibited CD8^+^ T cell proliferation (**Figure 2a**). These findings demonstrate that LDLR mediated LDL uptake is essential for CD8^+^ T cell clonal expansion.

**Figure 2.**
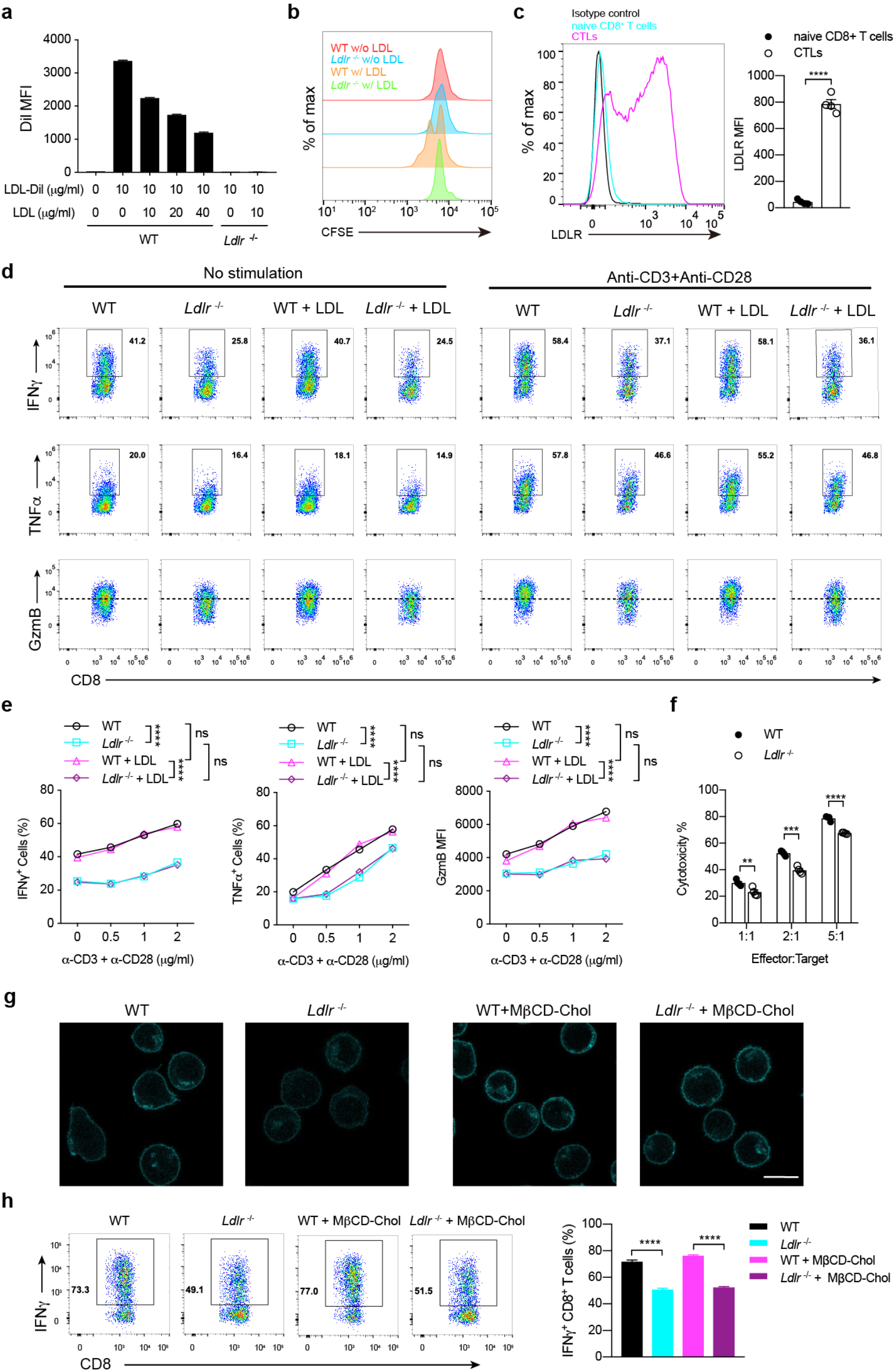
The regulation of LDLR on CD8^+^ T cell effector function is not fully dependent on LDL/cholesterol. **a**, LDL uptake of activated WT and *Ldlr*^-/-^ CD8^+^ T cells. CD8^+^ T cells were treated with LDL and LDL-Dil at indicated concentrations. The uptake of LDL-Dil was analyzed by flow cytometry. **b**, Proliferation of WT and *Ldlr*^-/-^ CD8^+^ T cells was measured by CFSE dilution, with or without the presence of LDL. **c**, LDLR expression was analyzed by flow cytometry of naïve CD8^+^ T cells and CTLs. Data were analyzed by *t* test (n = 4). **d, e**, Cytokine/granule productions of WT and *Ldlr*^-/-^ CTLs. CTLs were generated from the splenocytes of WT and *Ldlr*^-/-^ mouse and pretreated in LPDS medium for 2 hours, with or without the presence of LDL. The cells were then stimulated with anti-CD3 and anti-CD28 antibodies for 4 hours at indicated concentrations in corresponding medium. Data were analyzed by two-way ANOVA (n = 4). **f**, Cytotoxicity of WT and *Ldlr*^-/-^ CTLs. CTLs were pretreated with LPDS medium for 12 hours and cocultured with EL4 cells to determine the cytotoxicity. Data were analyzed by *t* test (n = 4). **g**, Filipin III staining to analyse cellular cholesterol distribution in untreated or MβCD-coated cholesterol treated WT and *Ldlr*^-/-^ CTLs. Scale bar, 10μm. **h**, IFNγ production of WT and *Ldlr*^-/-^ CTLs. Mature CTLs were generated from the splenocytes of WT and *Ldlr*^-/-^ mice and treated with MβCD-coated cholesterol or not. The cells were then stimulated with 1μg/ml plate-coated anti-CD3 and anti-CD28 antibodies for 4 hours. Data were analyzed by *t* test (n = 4). ns, no significance; **, *P* < 0.01; ***, *P* < 0.001; ****, *P* <0.0001. Error bars denote for the s.e.m.

CD8^+^ T cells experience cholesterol metabolic reprogramming when T-cell receptor (TCR) recognize the antigens. After which, the LDLR surface levels were dramatically increased in CTLs as compared with the naïve CD8^+^ cells (**Figure 2c**). To further evaluate the role of LDL in the effector function of CTLs, we re-stimulated the CTLs with anti-CD3 and anti-CD28 antibodies in medium supplemented with or without LDL. These results showed that a LDLR deficiency impaired CTL effector function, but that this effect could be lessened when supplemented with LDL (**Figure 2d, e**). CTL killing assay further verified this conclusion (**Figure 2f)**.

Cholesterol is the dominant component of LDL, and we found that the cholesterol levels of LDLR deficient CTLs were decreased, especially in the plasma membrane. Previous studies have demonstrated that the plasma membrane is involved in T cell activation (Gaus et al., 2005; Wu et al., 2016). To investigate whether LDLR-deficiency impaired CD8^+^ T cell effector function is reliant on plasma membrane cholesterol, we artificially increased the plasma membrane cholesterol levels of *Ldlr* knockout CTLs by adding MβCD-coated cholesterol, providing a cholesterol source independent of LDLR expression (**Figure 2g**). We then stimulated CTLs with anti-CD3 and anti-CD28 antibodies and evaluated cytokine production by flow cytometry. The results showed that increasing plasma membrane cholesterol did not improve the LDLR deficiency-induced effector function decline (**Figure 2h**). These data indicate that there is a mechanism by which LDLR regulates CD8^+^ T cell effector function, one which is not dependent on LDL or cholesterol.

### LDLR binds to TCR and regulates TCR signaling in CD8^+^ T cells

We then further investigated the underlying mechanisms by which LDLR regulates CD8^+^ T cell effector function. Notably, our results showed no significant differences in cytokine and granule production before antibody stimulation (**Figure 1c**), the defects of the *Ldlr* knockout appeared to be induced by anti-CD3 and anti-CD28 stimulation. Anti-CD3 and anti-CD28 stimulation mimics antigen recognition by the T-cell receptor (TCR) and the costimulatory signals by CD80/CD86-CD28 ligation, respectively (Trickett and Kwan, 2003). Previous studies have demonstrated that TCR mediated antigen recognition, or TCR signaling, is influenced by multiple factors, including kinases, phosphatases, and the plasma membrane lipid and protein composition (van der Merwe and Dushek, 2011; Stanford et al., 2012; Shi et al., 2013; Alcover et al., 2018). To evaluate the effect of LDLR deficiency on TCR signaling, we stimulated *Ldlr*^*-/-*^ CD8^+^ T cells with anti-CD3 and anti-CD28. We then detected the phosphorylation level of CD3ζ, a subunit of the TCR complex, and downstream signal pathways. The results showed that CD3ζ phosphorylation was inhibited by LDLR deficiency, as compared with the wild-type cells (**Figure 3a**). Consequently, the downstream signal pathways were also attenuated by LDLR deficiency (**Figure 3b**). Furthermore, the defects in TCR phosphorylation were not altered when we stripped cholesterol from the plasma membrane via MβCD treatment (**Figure 3c**). Together, these results suggest that LDLR may directly regulate TCR on the plasma membrane or the membrane proximal region.

**Figure 3.**
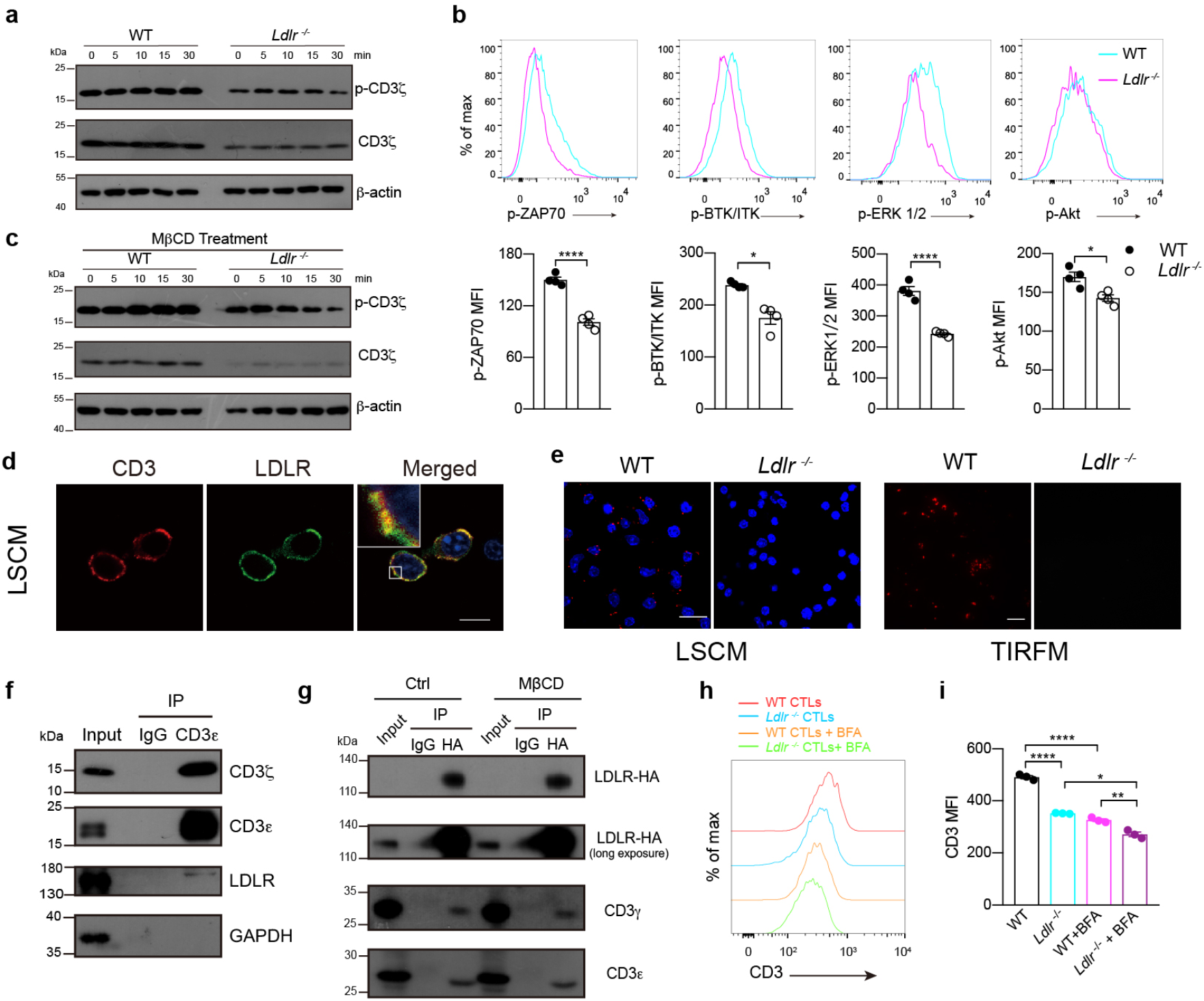
LDLR binds to TCR and regulate TCR signaling in CD8^+^ T cells. **a**, Immunoblotting to detect the phosphorylation of CD3 ζ of WT and *Ldlr*^-/-^ CTLs. CTLs were stimulated with 1μg/ml anti-CD3, anti-CD28, anti-Armenian hamster IgG and anti-Syrian hamster IgG for indicated times. **b**, Phosphorylation of ZAP70, BTK/ITK, ERK1/2 and Akt of WT and *Ldlr*^-/-^ CTLs. CTLs were stimulated as in (a) for 10 minutes. Data were analyzed by *t* test (n = 4).**c**, Immunoblotting to detect the phosphorylation of CD3ζ of MβCD treated WT and *Ldlr*^-/-^ CTLs. CTLs were stimulated as in (**a**).**d**, Fluorescence staining of CD3 and LDLR in CTLs. Scale bar, 10μm. LCSM, laser confocal scanning microscopy. **e**, Proximity Ligation Assay (PLA) analysis of CD3 and LDLR interaction in WT and *Ldlr*^-/-^ CTLs. Confocal images (left panel, scale bar, 20μm) and TIRFM images (right panel, scale bar, 10μm) were shown. Red, CD3-LDLR interaction signal; Blue, DAPI. TIRFM, total internal reflection fluorescence microscopy. **f**, CD3ε was immunoprecipitated (IP) in CTLs and its interaction with LDLR was analyzed by immunoblotting. **g**, HA-tagged LDLR was overexpressed in EL4 cells. The EL4 cells were treated with MβCD or not and then HA-tagged LDLR was immunoprecipitated with anti-HA antibody. The interaction between LDLR and CD3 was analyzed by immunoblotting. **h, i**, WT and *Ldlr*^-/-^ CTLs were treated with BFA (5μg/ml) or not for 2 hours. CD3 expression was analyzed by flow cytometry. Data were analyzed by *t* test (n = 3). *, *P* < 0.05; **, *P* < 0.01; ****, *P* <0.0001. Error bars denote for the s.e.m.

We next stained CD8^+^ T cells with anti-LDLR and anti-CD3 to determine the localization of these two proteins on the plasma membrane. Imaging data showed that LDLR colocalizes with CD3 on the plasma membrane of CD8^+^ T cells (**Figure 3d**). To further corroborate the interaction between LDLR and TCR complex, we used a PLA assay (Proximal Ligation Assay) to image the interaction. Confocal imaging data exhibited clear interaction spots in wild-type CD8^+^ T cells, but not in *Ldlr*^*-/-*^ CD8^+^ T cells (**Figure 3e**). Furthermore, TIRFM (Total Internal Reflection Fluorescence Microscopy) imaging showed the interaction of the LDLR and TCR complexes in the plasma membrane or the membrane proximal region of CD8^+^ T cells (**Figure 3e**). We then used a co-immunoprecipitation assay to determine the interaction between the CD3 subunit of TCR and LDLR. The results showed that there is indeed an interaction between the CD3 subunit and LDLR, and that this interaction is not influenced by the removal of plasma membrane cholesterol by MβCD (**Figure 3f, g**).

Additionally, we found that the surface TCR levels were reduced in *Ldlr*^-/-^ CD8^+^ T cells. To investigate further, we inhibited plasma membrane protein recycling via treatment with Brefeldin A and compared the surface TCR plasma membrane levels between *Ldlr*^-/-^ and wild-type CD8^+^ T cells (**Figure 3h, i**). These results thus suggest that LDLR may be involved in plasma membrane TCR recycling, thereby regulating TCR signaling and ultimately T cell effector function (**Supplementary figure 2**). Together, these experiments indicate that LDLR interacts with the TCR complex and regulates TCR signaling as an immune regulatory membrane protein, not just as a LDL transporter.

### Tumor-derived PCSK9 inhibits the antitumor activity of CD8^+^ T cells

To further investigate how the TME inhibits the LDLR level of tumor infiltrating CD8^+^ T cells, we transferred OT-I CTLs to *Rag2*^-/-^ mice with MC38-OVA tumors. Then, we isolated the tumor infiltrating antigen specific CD8^+^ T cells and quantified the mRNA levels and cell surface expression of LDLR by qPCR and flow cytometry, respectively. The results show that cell surface LDLR levels were dramatically decreased during early stage T cell infiltration, while conversely, the mRNA levels remained normal (**Figure 4a, b**). This finding indicates that there is another pathway that regulates the cell surface LDLR levels besides transcriptional regulation.

**Figure 4.**
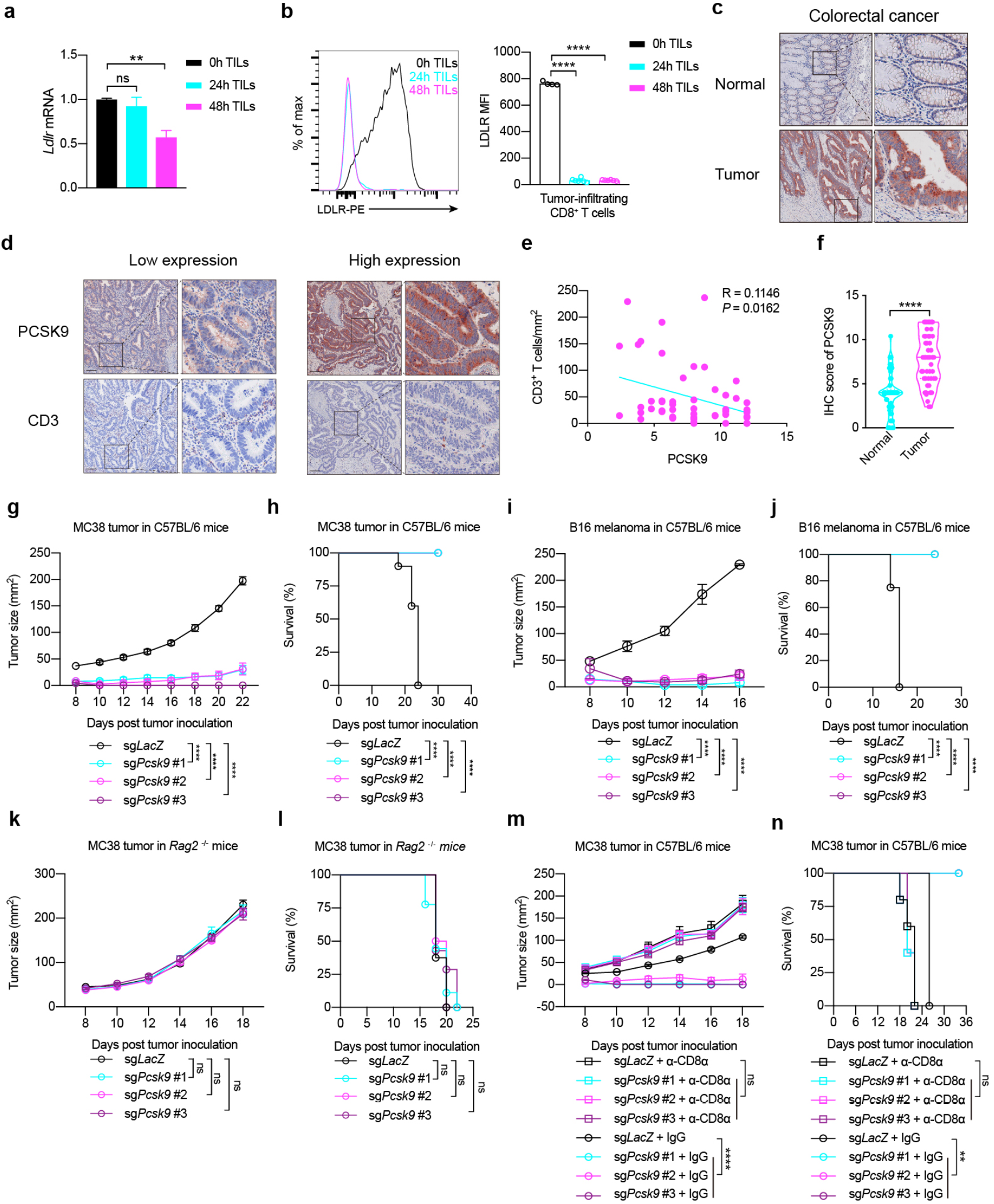
Tumor-derived PCSK9 inhibits the antitumor activity of CD8^+^ T cells. **a**, LDLR expression was assessed in tumor infiltrating CD8^+^ T cells (TILs) at 0, 24 or 48 hours post adoptive transfer. Data were analyzed by *t* test (n = 4-6). **b, c**, Human normal or tumor colorectal sections were stained with anti-PCSK9 antibody by immunohistochemistry and the abundance of PCSK9 was assessed in (c). Data were analyzed by *t* test (n = 50). **d, e**, PCSK9 and CD3 staining were shown in PCSK9 low-expression and high-expression tumors. Pearson correlation coefficient (R) and *P* value (*P*) of PCSK9 expression and CD3^+^ cells infiltration were analyzed in (e). **f, g**, Tumor growth (f) and survival (g) of *Pcsk9* knockout MC38 tumor-bearing C57BL/6 mice. Data were analyzed by two-way ANOVA (n = 10). **h, i**, Tumor growth (h) and survival (i) of *Pcsk9* knockout B16F10 melanoma-bearing C57BL/6 mice. Data were analyzed by two-way ANOVA (n = 7-8). **j, k**, Tumor growth (j) and survival (k) of *Pcsk9* knockout MC38 tumor-bearing Rag2^-/-^ mice. Data were analyzed by two-way ANOVA (n = 7-9). **l, m**, Tumor growth (l) and survival (m) of *Pcsk9* knockout MC38 tumor-bearing C57BL/6 mice with CD8α^+^ cells depletion. Data were analyzed by two-way ANOVA (n = 5-6). Scale bar, 120μm. ns, no significance; **, *P* < 0.01; ****, *P* <0.0001. Error bars denote for the s.e.m.

PCSK9, a previously identified LDLR modulator and a therapeutic drug target for treating hypercholesterolemia, has been implicated in a critical role for regulating LDLR protein levels via mediating LDLR internalization and degradation (Abifadel et al., 2003; Maxwell et al., 2005; Cunningham et al., 2007; Zhang et al., 2007; Liu et al., 2020). To determine PCSK9 involvement in surface LDLR regulation, we first collected clinical samples of colorectal cancer (CRC) and detected PCSK9 expression by immunohistochemistry (IHC). IHC scoring showed there was higher PCSK9 expression in cancerous regions than in the adjacent normal region (**Figure 4c, f**). Furthermore, when we detected the CD3 levels in the CRC samples, we found there was a significant negative correlation between CD3^+^ T cell infiltration and PCSK9 level (**Figure 4d, e**).

To further evaluate the relationship of T cell infiltration and PCSK9 expression in tumors, we depleted *Pcsk9* gene expression in a mouse CRC cell line (MC38) and melanoma cell line (B16F10) via CRISPR/Cas9. We then transplanted the gene modified tumor cells into wild-type syngeneic mice. The results showed that PCSK9 depletion inhibited tumor progression and greatly extended mice survival time (**Figure 4g-j**). Conversely, when we transplanted the MC38 tumor cells to *Rag2*^-/-^mice, exhibiting T cell and B cell deficiency, we found there were no significant differences between the wild-type MC38 and *Pcsk9* knockout MC38 mice (**Figure 4k, l**).A similar result was achieved by using shRNA to induce*Pcsk9* knockdown in MC38 tumor (**Supplementary figure 3a-e**), where the tumor infiltrating CD8^+^ T cells in *Pcsk9*-knockdown tumors showed increased antitumor activity (**Supplementary figure 3f**). These finding indicate that the lower progression of *Pcsk9* knockout tumor in immunocompetent mice may be attributed to the antitumor activity of adoptive immune cells, like T cells and B cells. Given that CD8^+^ T cells play critical roles in antitumor immunity, we used anti-CD8 antibody to deplete CD8^+^ T cells *in vivo*, to determine on which cell type the impact of PCSK9 was most prominent. Our data showed that when CD8^+^ T cells were depleted, there were no significant differences between the wild-type MC38 and the *Pcsk9*-knockout MC38 tumors in the syngeneic immunocompetent mice (**Figure 4m, n**). Collectively, these results demonstrate that the tumor derived PCSK9 predominantly inhibits the immune response of CD8^+^ T cells in achieving immune evasion.

Notably, we also investigated the intrinsic effect of PCSK9 on CD8^+^ T cells. The syngeneic mouse tumor model showed that tumor progression was inhibited in *Pcsk9*^-/-^ mice when compared with the wild-type mice. We further stimulated the splenic naïve CD8^+^ T cells from *Pcsk9*^-/-^ mice with anti-CD3 and anti-CD28 antibodies to detect cytokine and granule production. The results showed PCSK9-deficent CD8^+^ T cells exhibited higher effector function. Moreover, we found that PCSK9 intrinsically inhibited CD8^+^ T cell function through evaluating the immunological synapse formation and cytotoxicity as well as antitumor activity *in vivo* through adoptive T cell transfer assay. These findings indicate that PCSK9 intrinsically inhibits the antitumor activity of CD8^+^ T cells.

### PCSK9 inhibits CD8^+^ T cell antitumor activity via LDLR and TCR signaling inhibition

To further investigate the mechanisms behind how PCSK9 regulates CD8^+^ Tcell antitumor activity, we transplanted wild-type MC38-OVA or *Pcsk9*-depleted MC38-OVA cells into *Rag2*^-/-^ mice. We then transferred wild-type OT-I CTLs or *Ldlr*^-/-^ OT-I CTLs into the tumor burdened mice. The results showed that there was no significant difference in tumor progression between the wild-type MC38 tumor and *Pcsk9*-depleted MC38 tumor in *Rag2*^-/-^ mice who did not receive the CTL transfer (**Figure 5a**). In accord with our earlier findings, the antitumor activity of the wild-type CTLs was higher in the *Pcsk9*-depleted MC38 tumor than in the wild-type MC38 tumor (**Figure 5b**). Conversely, when we transferred *Ldlr*^-/-^ OT-I CTLs into the mice, there were no significant differences in tumor progression between the wild-type MC38 tumor and the *Pcsk9*-depleted MC38 tumor (**Figure 5c**). These findings indicate that the PCSK9-derived inhibition of CD8^+^ T cell’s antitumor activity is through LDLR. Concurrently, we treated CD8^+^ T cells with recombinant mouse PCSK9 protein.The results showed that the surface level of LDLR in CD8^+^ T cells was reduced by PCSK9 treatment and consequently, the plasma membrane TCR level, CD3 phosphorylation, and the effector function were all down regulated (**Figure 5d-g**). Furthermore, we found that PCSK9 treatment inhibited cytokine production of CD8^+^ T cells when we pretreated CD8^+^ T cells with recombinant mouse PCSK9. We then used the *in vitro* killing assay to assess the influence of PCSK9 on CTL cytotoxicity, with PCSK9 over-expressing EL4 cells—which show normal MHC I (H2K^b^) expression—as the target cells. We found that the overexpression of PCSK9 substantially inhibited the killing efficiency of OT-I CD8^+^ T cells (**Figure 5h**), findings which are consistent with the conclusion from LDLR deficient CD8^+^ T cells.

**Figure 5.**
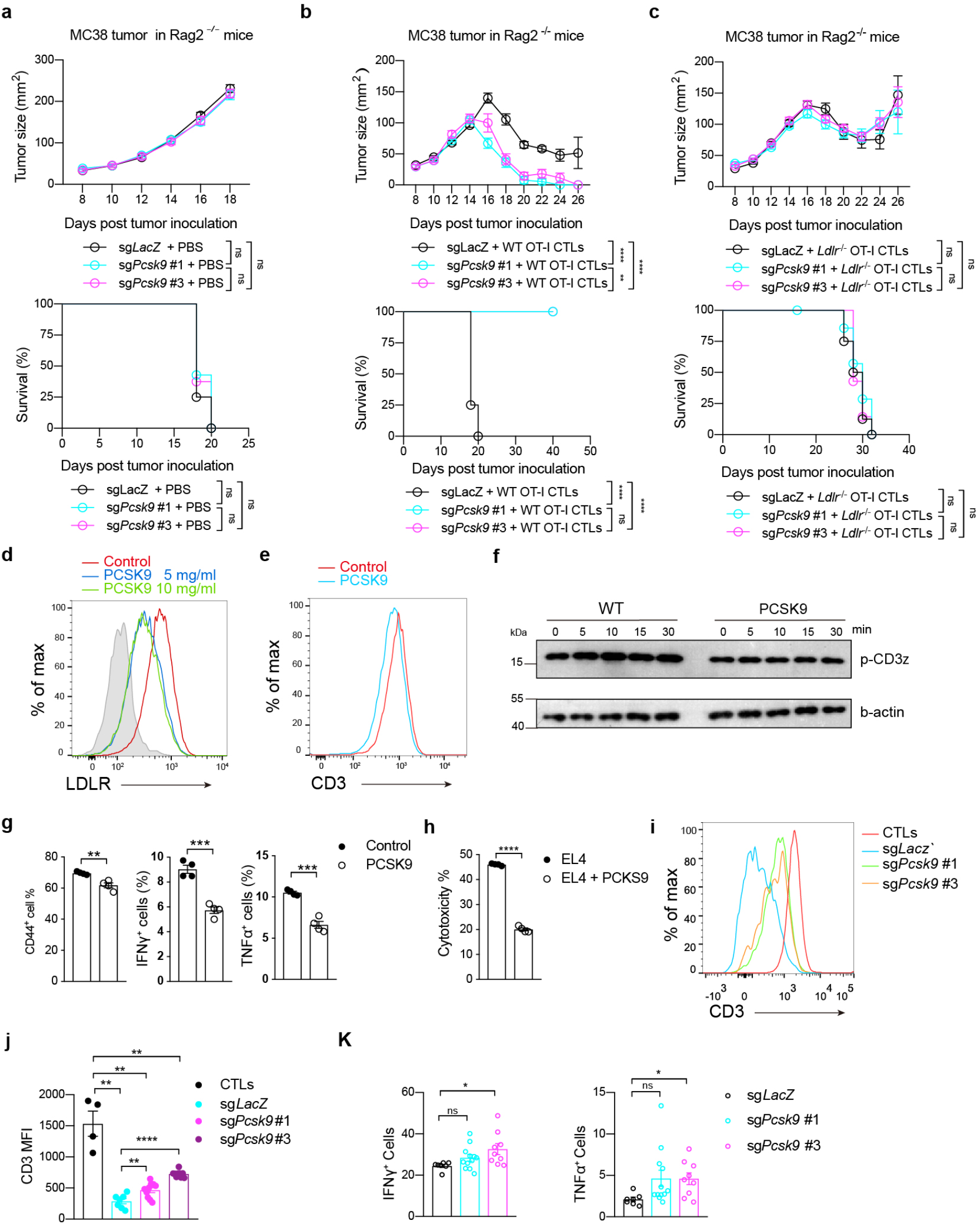
PCSK9 inhibits CD8^+^ T cell antitumor activity via LDLR and TCR signaling inhibition a-c, Tumor growth and survival of *Pcsk9* knockout MC38-OVA tumor-bearing *Rag2*^-/-^ mice after adoptive transfer of PBS (a), WT CTLs (b) or *Ldlr*^-/-^ CTLs (c). Data were analyzed by two-way ANOVA (n=7-8). **d**, LDLR expression was measured in PCSK9-treated CTLs by flow cytometry. CTLs were treated with PCSK9 protein at indicated concentrations for 6 hours. **e**, CTLs were treated with 5μg/ml PCSK9 protein for 6 hours. CD3 expression was measured by flow cytometry. **f**, Immunoblotting to detect the phosphorylation of CD3ζ in control and PCSK9-treated CTLs. CTLs were pretreated with 5μg/ml PCSK9 protein for 6 hours and stimulated with 1μg/ml anti-CD3, anti-CD28, anti-Armenian hamster IgG and anti-Syrian hamster IgG for indicated times. **g**, T cell activation and cytokine productions of PCSK9 treated activated CD8+ T cells. Naïve CD8+ T cells were isolated and stimulated with 2μg/ml anti-CD3 and anti-CD28 in the presence or absence of PCSK9 protein (5 μg/ml). Data were analyzed by *t* test (n=4).**h**, Cytotoxicity of WT CTLs cocultured with PCSK9 overexpressed EL4 cells. PCSK9 was overexpressed in EL4 cells by retrovirus infection. CTLs were cocultured with the EL4 cells to determine the cytotoxicity. Data were analyzed by *t* test (n=4). **i, j**, CD3 surface levels were analyzed by flow cytometry in CTLs and TILs isolated from *Pcsk9* knockout MC38-OVA tumors at Day7 post adoptive transfer. Data were analyzed by *t* test (CTLs, n = 4; TILs, n = 7-10). **l**, IFNγ production in isolated TILs of *Pcsk9* knockout MC38-OVA tumors. ns, no significance; *, *P* < 0.05; **, *P* < 0.01; ***, p<0.001; ****, *P* <0.0001. Error bars denote for the s.e.m.

To further evaluate the *in vivo* effects of PCSK9 on TCR, we transplanted wild-type and *Pcsk9*-depleted MC38-OVA cells to *Rag2*^-/-^ mice, respectively. Then, we transferred OT-I CTLs to the tumor burdened mice. At day 7 post tumor inoculation, we isolated the tumor infiltrating CD8^+^ T cells and performed flow cytometric analysis. The results showed that the TME inhibited the levels of surface TCR and the effector function but that PCSK9 depletion alleviated this inhibition (**Figure 5i-k**), the. Collectively, these data demonstrated that the tumor derived PCSK9 may down regulate LDLR and TCR signaling and effector function of CD8^+^ T cells, thus inhibiting the antitumor activity of CD8^+^ T cells in the TME.

### Inhibiting PCSK9 potentiates the antitumor activity of CD8^+^ T cells

Targeting the PCSK9/LDLR axis has shown clinical success in treating hypercholesterolemia and multiple drugs, such as evolocumab and alirocumab, have been approved for clinical use. Herein, we intensively investigated the PCSK9/LDLR axis in the CD8^+^ T cell antitumor immune response. To evaluate whether targeting the PCSK9/LDLR axis possesses clinical cancer treatment potential, we used syngeneic mouse models to determine the antitumor effect of PCSK9 inhibitors. The blocking antibodies used, evolocumab and alirocumab, were humanized antibodies. Previous research has found that the binding affinity of evolocumab to mouse PCSK9 (Kd=17 nM) is 1000-fold less than its binding affinity to human PCSK9 (Kd=16 pM) (Brody and Brody, 2018). Similarly the binding affinity of alirocumab to mouse PCSK9 (Kd=2.61 nM) is 4.5 fold less than the binding affinity to human PCSK9 (Kd=0.58 nM). *In vitro* experiment also demonstrated that alirocumab is substantially less effective on mouse PCSK9 compared to human PCSK9. Therefore, we used a chemical inhibitor, PF0644684,which has been demonstrated previously to effectively inhibit PCSK9 expression through slowing down PCSK9 translation (Lintner et al., 2017) .

First, we assessed the inhibitory effect of PF0644684 on tumor PCSK9 *in vivo* and found that 8 administrations of a 2-10 mg/kg dose and effectively inhibited PCSK9 expression in MC38 tumors in C57BL/6 mice (**Supplementary figure 5a-b**). We then further evaluated the antitumor effect of PF0644684 in the syngeneic mouse tumor model, including MC38 and B16 tumors, where administration of PF0644684 effectively inhibited tumor progression (**Figure 6a-d**). In contrast, there was no analogous antitumor effect with PF0644686 administration in MC38 tumor burdened *Rag2*^-/-^ mice, which lack CD8^+^ T cells (**Figure 6e, f**). These findings were consistent with those of the *Pcsk9*^-/-^ tumor cells. Furthermore, the *in vitro* CTL killing assay showed that EL4-OVA cells pretreated with PF0644684 increased the cytotoxicity of OT-I CTLs to the target cells (**Supplementary figure 5c**). Collectively, these findings indicate that PCSK9 inhibition potentiates the antitumor activity of CD8^+^ T cells.

**Figure 6.**
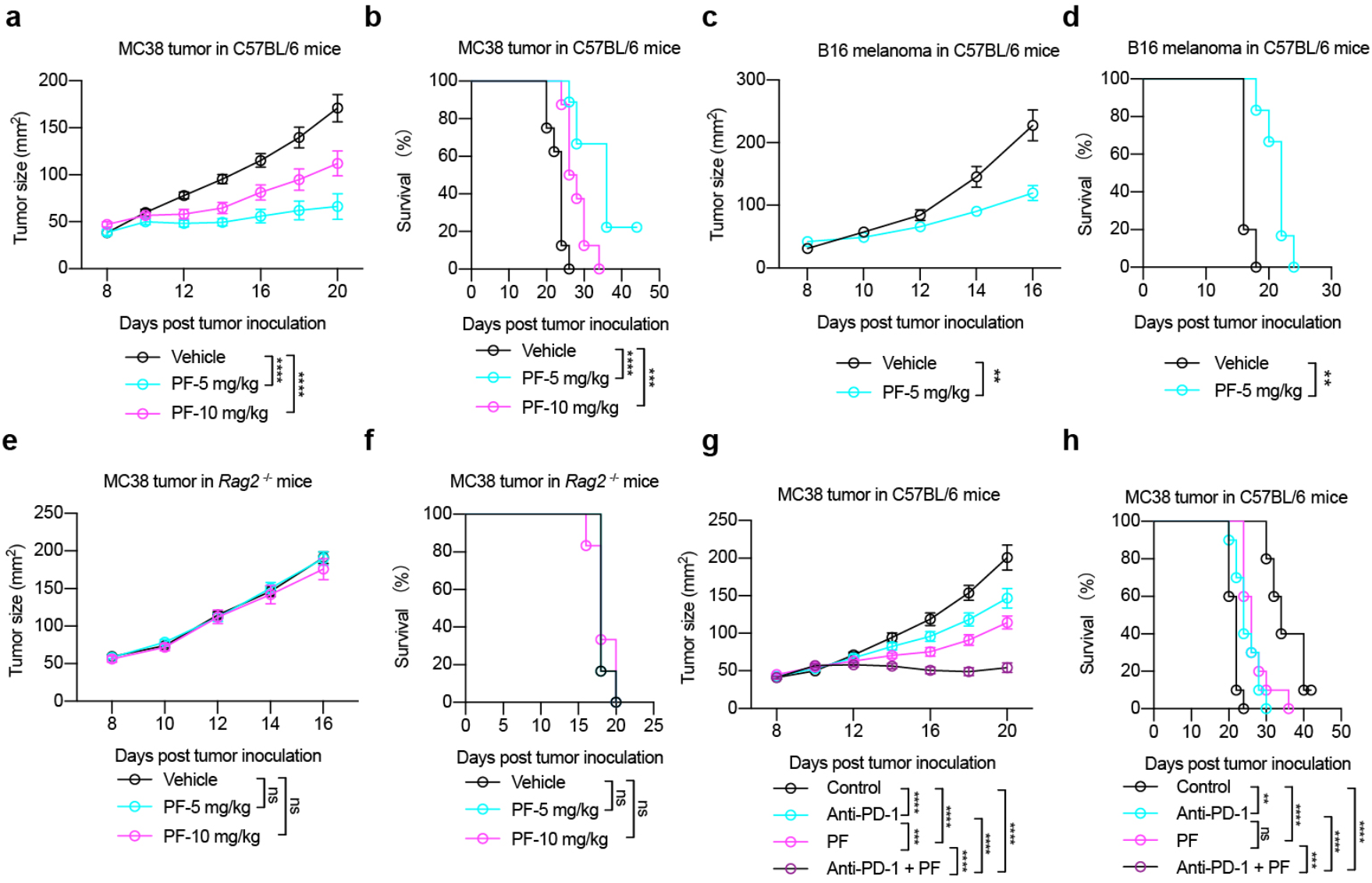
Inhibiting PCSK9 potentiating the antitumor activity of CD8^+^ T cells. **a, b**, Tumor growth (a) and survival (b) of MC38 tumor-bearing C57BL/6 mice. Vehicle, 5mg/kg or 10mg/kg PF0644684 were injected intraperitoneally every 2 days. Data were analyzed by two-way ANOVA (n = 8-9). **c, d**, Tumor growth (c) and survival (d) of B16F10 melanoma-bearing C57BL/6 mice. Vehicle or 5mg/kg PF0644684 were injected intraperitoneally every 2 days. Data were analyzed by two-way ANOVA (n = 8-9). **e, f**, Tumor growth (e) and survival (f) of MC38 cells on Rag2^-/-^ mice. PF0644684 was injected intraperitoneally as in (a, b). Data were analyzed by two-way ANOVA (n = 6). **g, h**, A combined therapy (PF0644684 and anti-PD-1) or monotherapies (PF0644684 or anti-PD-1) in treating MC38 tumors on C57BL/6 mice. Tumor growth (g) and survival (h) were shown. Data were analyzed by two-way ANOVA (n = 10). ns, no significance; **, *P* < 0.01; ***, *P* < 0.001; ****, *P* <0.0001. Error bars denote for the s.e.m.

We then tested a combination therapy of PCSK9 inhibition and immune checkpoint blockade therapy to observe potential synergistic impacts. We treated MC38 tumor burdened C57BL/6 mice, which are immunocompetent syngeneic mice, with PF0644684 and anti-PD1 antibodies (**Figure 6g, h**). The results showed that the combination therapy had a stronger tumor suppressive effect than either monotherapy, highlighting that PCSK9 inhibition has potential as a novel cancer immunotherapy strategy.

## DISCUSSION

T cells undergo distinctive metabolic reprograming in different stages, and these metabolic regulations have been demonstrated to play critical roles in T cells’ immune responses (Ecker et al., 2018; Geltink et al., 2018; Chapman et al., 2020). As a main component of cellular metabolism, cholesterol metabolism is essential for effective T cell immune responses. But precisely how cholesterol metabolic pathways regulate CD8^+^ T cell function and how metabolic reprogramming regulates CD8^+^ T cell antitumor activity, needs more extensive and comprehensive investigation. Our previous study, and several related studies, have shown that the storage and biosynthetic pathwaysof cholesterol play an important role in the regulation of the CD8^+^ T cell immune response (Bensinger et al., 2008; Yang et al., 2016; Ma et al., 2018; Ma et al., 2019). These studies support that CD8^+^ T cells need free cholesterol to support effector function and clonal expansion. The tumor microenvironment has been demonstrated as a hypoxia, nutrient restrictedenvironment (Zhang and Ertl, 2016). Can CD8^+^ T cells obtain sufficient cholesterol in the tumor microenvironment to support their effector function and antitumor activity? And if so how? To answer thesequestions, we measured the cholesterol/LDL distribution in cancerous and paracancerous normal tissues in mice models and clinical sample from cancer patients. We found that APOB, which is a marker of LDL, showed higher levels in tumor regions compared with normal tissues. However, the cell cholesterol level of tumor infiltrating CD8^+^ T cells were substantially lower than those of peripheral CD8^+^ T cells, suggesting that the cholesterol metabolic pathways might be reprogramed. Further study confirmed this hypothesis, the cholesterol biosynthesis pathway and up take pathway by LDLR were found to be suppressed in the tumor microenvironment.

LDLR has been previously well characterized as a transporter of LDL, andLDLR deficiency has been identified asthe cause of high serum LDL, hypercholesterolemia, and other related metabolic dysfunction diseases (Hobbs et al., 1990; Bayes-Genis et al., 2017; Da Dalt et al., 2019). The downregulation of LDLR might be a significant factor influencing the cellular cholesterol levels of tumor infiltrating CD8^+^ T cells. Our *in vitro* and *in vivo* experiment demonstrated that LDLR is in fact necessary for CD8^+^ T cell antitumor immunity. When we assessedthe function of LDL/cholesterol in CD8^+^ T cells, we found LDL/cholesterol is essential for CD8^+^ T cell proliferation, but not effector function, particularlyin activated cytotoxic CD8^+^ T cells (CTLs). Moreover, we found that LDLR interacts with the T-cell receptor (TCR) on the plasma membrane of CD8^+^ T cells. This interaction favors TCR signaling and the effector function of CD8^+^ T cells. LDLR deficiency appears to inhibit TCR recycling to the plasma membrane as well as TCR signaling. Taken together, we found a noncanonical function of LDLR, in which it functions as a membrane protein to regulate the other receptors on the plasma membrane, not just as a LDL/cholesterol transporter. This finding indicates that LDLR could regulated other membrane proteins and may be involved in more physiological functions in different cell types, highlighting it as a candidate for further study.

After elucidating the critical role of LDLR, the next question was how does the tumor microenvironment inhibit LDLR expression in CD8^+^ T cells? Generally, protein expression can be inhibited at two levels: the transcriptional level and the protein level. In T cells, LDLR transcription is indirectly regulated by TCR/CD28 signaling. T cell activation by antigen stimulation can up regulate LDLR mRNA level (Yang et al., 2016). Moreover, T cell activation may downregulate IDOL, which is the E3 ligase of LDLR and mediates LDLR ubiquitination and degradation (Zelcer et al., 2009; Yang et al., 2016). In the past years, PCSK9, which has been shown to be a negative modulator of LDLR, has been utilized as a clinical drug target for treating hypercholesterolemia (Stein et al., 2013; Raal et al., 2015; Raal et al., 2017). We found that PCSK9 was highly expressed in the tumor region of CRC patients and that T cell infiltration was negatively correlated with the PCSK9 levels. Our findings suggest that tumor cell derived PCSK9 may downregulate the surface LDLR level in CD8^+^ T cells, thereby inhibiting the antitumor activity of CD8^+^ T cells. Given that the LDLR levels of CD8^+^ T cells were downregulated during early stage infiltration (12-24hours, Figure 1B)—at which time the transcription of *Ldlr* was not altered—and in combination with the finding that LDLR may directly regulated TCR signaling—which is necessary for *Ldlr* transcription—we speculate that the tumor microenvironment derived PCSK9 may be the source of LDLR downregulation and consequently, the immune suppression of CD8^+^ T cells. This speculation was confirmed in the *in vivo* syngeneic mouse tumor model, where the depletion of tumor PCSK9 alleviated the immune suppression by the tumor microenvironment on CD8^+^ T cells. Moreover, when we examined the intrinsic function of PCSK9 in CD8^+^ T cells, we found that PCSK9 intrinsically inhibited the effector function of CD8^+^ T cells, with the PCSK9 knockout CD8^+^T cells exhibiting higher antitumor activities. Which, indicates that the simultaneous inhibition of PCSK9 expression intumors and CD8^+^ T cells maybe a therapeutic approach to potentiate CD8^+^ T cell antitumor immunity.

Targeting metabolic reprogramming has been demonstrated as a potential method for cancer immunotherapy (Dugnani et al., 2017; Kishton et al., 2017; Sukumar et al., 2017). To further assess theclinical potential of inhibiting PCSK9, we used a chemical inhibitor of PCSK9, PF0644684, which has proven to inhibit PCSK9 translation (Lintner et al., 2017). This inhibitor successfully demonstrated antitumor activity in a syngeneic mouse tumor model and when used in combination with anti-PD-1 antibodies, the antitumor effect was further enhanced. These finding further support that targeting the metabolic pathway of cholesterol is a potential method for cancer immunotherapy. In summary, we have demonstrated that LDLR functions as a critical immune regulatory receptor for CD8^+^ T cells in the tumor microenvironment. Furthermore, we report a novel mechanism for LDLR activity, whereby it interacts with TCR and regulates TCR signaling, ultimately impacting CD8^+^ T cells effector function. Further investigation revealed that tumor derived PCSK9 is the critical factor for immune suppression of CD8^+^ T cells by the tumor microenvironment. Collectively, our findings highlight that the PCSK9-LDLR axis is the metabolic immune checkpoint of the tumor microenvironment and that targeting this pathway holds great potential in cancer immunotherapy.

## Methods and materials

### Patients and clinical specimens

The paraffin embedded tissues of colorectal carcinoma (CRC) tissues, adjacent non-carcinoma tissues (ANT), lung cancer tissues and normal lung tissues were obtained from the tissue bank of the Department of Pathology, Nanfang Hospital, Southern Medical University. Samples were collected from colorectal cancer, lung cancer and breast cancer that had been clinically diagnosed as cancer. The study protocols concerning human subjects are consistent with the principles of the declaration of Helsinki. The study was approved by the Clinical Research Ethics Committee of Southern Medical University.

### Mice

C57BL/6 mice, *Rag2*^*–/–*^mice, *Ldlr*^*–/–*^ mice, *PCSK9*^*–/–*^ mice and OT-I TCR transgenic mice were originally purchased from the Jackson Laboratory. Through mouse crossing, *Ldlr*^*–/–*^ OT-I mice and *PCSK9*^*–/–*^ OT-I mice were obtained and the genotypes were validated by using PCR. All mice used in this study are maintained in specific pathogen-free conditions. All animal experiments used mice were randomly allocated to specific groups with matched age and sex. All animal experiments were approved by the Ethics Committee on Use and Care of Animals of Southern Medical University.

### Reagents and antibodies

For flow cytometric analysis, anti-CD3ε (145-2C11), anti-CD8 (53-6.7), anti-CD44 (IM7), anti-CD45 (30-F11), anti-IFNγ(XMG1.2), anti-granzyme B (NGZB), anti-TNF-α (MP6-XT22), anti-p-ZAP70/Syk(Tyr319, Tyr352) (n3kobu5), anti-p-BTK/ITK(Tyr551, Tyr511) (M4G3LN), anti-p-Akt (Ser473) (SDRNR) and anti-p-Erk (Thr202, Tyr204) (MILAN8R) were purchased from Thermofisher. Anti-mouse LDLR (101) was purchased from Sino Biological Inc. For western blot analysis, anti-β-actin, anti-GAPDH, anti-CD3ε, anti-CD3γ, anti-CD3ζ were from Santa Cruz Biotechnology. Anti-p-CD3ζ (Tyr142) was from Abcam. Anti-HA was from Sigma. For immunohistochemistry analysis, anti-Apolipoprotein B (Abcam), anti-PCSK9 (Sino Biological) and anti-CD3 (SP7, Abcam) were purchased from indicated companies. For immunofluorescence and PLA staining, anti-LDLR was from Lifespan. Anti-CD3 was from Genetex. Anti-CD3ε was from Bio X Cell. Filipin III was from Cayman. PF-06446846 was from MedChemExpress. For tissue infiltrated T cells isolation, Type IV Collagenase was from Gibco. DNase I was from Applichem. Hyaluronidase was from Sigma. Percoll was from GE. Anti-CD3ε(145-2C11, Bio X Cell), anti-human CD3(UCHT1, Bio X Cell) anti-mouse CD28(37.51, Bio X Cell) and anti-human CD28(9.3, Bio X Cell) were used for T cell activation. OVA257-264 peptide (SIINFEKL) was from ChinaPeptides Co.. PCSK9 protein was purchased from ACROBiosystems. Celltrace CFSE, Celltracker Deep Red and Cell proliferation Dye eFluor 450 were from Invitrogen. Methyl-beta-cyclodextrin (MβCD) and MβCD-coated cholesterol were purchased from Sigma.

### Cell lines

MC38 cells were provided by JENNIO Biological Technology (Guangzhou, China). B16F10 and EL-4 cells were originally obtained from the American Type Culture Collection (ATCC), and proved mycoplasma-free. MC38, B16F10 and 293T cells were maintained in DMEM (Gibco) and EL-4 cells were in RPMI-1640 (Gibco) medium respectively, supplemented with 10% FBS and 1% penicillin-streptomycin. Cells were cultured at 37°C in a humidified atmosphere containing 95% air and 5% CO2. MC38-OVA and B16F10-OVA cells were generated by lentivirus infection and mCherry^+^ cells were sorted as OVA^+^ cells.

### PCSK9 knockdown and knockout cell lines

To generate PCSK9 knockdown cell lines, lentiviruses were produced by transfecting 293T cells with pLKO.1-GFP, psPAX2 and VSV-G plasmids. MC38 cells were infected with pLKO.1 shRNA lentivirus and GFP^+^ cells were selected by Fluorescence-activated Cell Sorting. Knockdown efficiency was determined by QPCR. ShRNA sequences against PCSK9 were as follows: sh*Pcsk9* #1: 5’-GCTGATCCACTTCTCTACC-3’; sh*Pcsk9* #2: 5’-CAGAGGCTACAGATTGAAC -3’.

To generate PCSK9 knockout cells, lentiviruses were produced by transfecting 293T cells with Lenti-CRISPR-V2, psPAX2 and VSV-G plasmids. MC38 and B16F10 cells were infected with lentivirus and GFP^+^ cells were selected by Fluorescence-activated Cell Sorting. To generate PCSK9 knockout single cell clones, the cells were digested, limited diluted and finally plated on 96-well plates at a concentration of 0.8 cell per well, which was confirmed visually. Wells containing either none or more than one cell were excluded for further analysis. The genotypes of single cell clones were identified by Sanger sequencing. SgRNA sequences targeting mouse PCSK9 were as follows: sg*Pcsk9* #1, 5’-GCTGATGAGGCCGCACATG-3’; sg*Pcsk9* #2, 5’-CTACTGTGCCCCACCGGCGC-3’; sg*Pcsk9* #3, 5’-ACTTCAACAGCGTGCCGG -3’, SgRNA sequence targeting LacZ: 5’-GCGAATACGCCCACGCGAT-3’.

### Flow cytometic analysis

Anti-mouse CD16/32 antibody was used to block non-specific binding to Fc receptors before all surface staining. For surface staining, cells were collected and staining with antibodies at 4°C for 30 min. For cytokine staining, cells were stimulated with Brefeldin A (3 µg/ml, invitrogen) for 4 hours before cells were harvested for analysis. Before intracellular staining and phosphorylation staining, harvested cells were stained the surface protein and then fixed with 4% PFA for 5 minutes at RT. Then the cells were permeabilized with 0.1% Triton X-100 for 5 minutes at RT. Then the cells were stained with specific antibodies for 1 hour at 4°C. Flow cytometric data were analyzed with a SONY SA3800 flow cytometer and FlowJo software (Treestar).

### Immunohistochemistry

Human tissue samples and mouse tumor tissues were embedded with paraffin and sectioned longitudinally at 5 µm. All tissue sections were de-waxed and rehydrated and then antigens were retrieved with 10 mM sodium citrate (pH 6.0) in a pressure cooker. Incubated sections in 0.3% H2O2 in methanal for 30 min for blocking endogenous peroxidase activity. The slides were blocked with goat serum and then processed for against human or mouse PCSK9, human ApoB and human CD3 at 4°C overnight. Then the slides were incubated with a goat anti-HRP IgG antibody and developed with 3-amino-9-ethylcarbazole (ACE) and counterstained with hematoxylin. Images were captured by use of Zeiss microscope. Immunohistochemical results were scored in accordance with immunoreactive score (IRS) standards proposed by Remmele and Stegner. IRS = SI (staining intensity) × PP (percentage of positive cells). Negative PP, 0; 10% PP, 1; 10-50% PP, 2; 51-80% PP, 3; and > 80% PP, 4. Negative SI, 0; Mild SI, 1; Moderate SI, 2; Strongly positive SI, 3. Images were scored independently by two pathologists who were blinded to patient information.

### Real time RT-PCR

Total RNA was extracted with TRIzol reagent (Thermofisher). cDNA was synthesized with the Hiscript III RT Supermix for qPCR Kit (Vazyme) according to the manufacturer’s instructions. Real-time quantitative PCR using gene specific primers (5’-3’): *18s* (forward, TTGATTAAGTCCCTGCCCTTTGT; reverse, CGATCCGAGGGCCTCACTA); *Ldlr* (forward, TGACTCAGACGAACAAGGCTG, reverse, ATCTAGGCAATCTCGGTCTCC); *Srebp1* (forward, GCAGCCACCATCTAGCCTG; reverse, CAGCAGTGAGTCTGCCTTGAT); *Srebp2* (forward, GCAGCAACGGGACCATTCT; reverse, CCCCATGACTAAGTCCTTCAACT); *Acaca* (forward, ATGGGCGGAATGGTCTCTTTC; reverse, TGGGGACCTTGTCTTCATCAT); *Fasn* (forward, GGAGGTGGTGATAGCCGGTAT; reverse, TGGGTAATCCATAGAGCCCAG); *Hmgcs* (forward, AACTGGTGCAGAAATCTCTAGC; reverse, GGTTGAATAGCTCAGAACTAGCC); *Hmgcr* (forward, AGCTTGCCCGAATTGTATGTG; reverse, TCTGTTGTGAACCATGTGACTTC); *Sqle* (forward, ATAAGAAATGCGGGGATGTCAC; reverse, ATATCCGAGAAGGCAGCGAAC); *Idol* (forward, TGCAGGCGTCTAGGGATCAT; reverse, GTTTAAGGCGGTAAGGTGCCA); *Abca1* (forward, AAAACCGCAGACATCCTTCAG; reverse, CATACCGAAACTCGTTCACCC); *Abcg1* (forward, CTTTCCTACTCTGTACCCGAGG; reverse, CGGGGCATTCCATTGATAAGG); *Acat1* (forward, GAAACCGGCTGTCAAAATCTGG; reverse, TGTGACCATTTCTGTATGTGTCC); *Acat2* (forward, ACAAGACAGACCTCTTCCCTC; reverse, ATGGTTCGGAAATGTTCACC); *Nceh* (forward, TTGAATACAGGCTAGTCCCACA; reverse, CAACGTAGGTAAACTGTTGTCCC); *Ifng* (forward, ATGAACGCTACACACTGCATC; reverse, CCATCCTTTTGCCAGTTCCTC); *Pcsk9*(forward, GAGACCCAGAGGCTACAGATT; reverse, AATGTACTCCACATGGGGCAA). All PCR reactions were conducted on a QuantStudio real-time PCR system (Thermo Fisher) in triplicates. Gene expression was normalized to 18s.

## CD8^+^ T cell isolation and activation

Naïve CD8^+^ T cells were isolated from mouse spleen by a EasySep Mouse Naïve CD8^+^ T cell Isolation Kit (Stem Cell). Then the cells were stimulated with plate-coated anti-CD3 and anti-CD28 at indicated concentration for indicated times.

### CTL generation

OT-I mouse splenocytes were harvested and homogenized using sterile techniques. Red blood cells were then lysed with ACK buffer for 5 min at RT. The splenocytes were pelleted and resuspended at 1 × 10^6^ per millilitre in RPMI-1640 medium with 10% FBS, 1% penicillin-streptomycin, 2-mercaptoethanol and supplemented with 10 nM OVA257-264 peptide and 10 ng/ml human recombinant interleukin-2 (Peprotech) for 3 days. Then the cells were cultured in fresh medium containing 10 ng/ml IL-2 for 2 more days to do the subsequent experiments.

### Measurement of CD8 T-cell proliferation

Isolated naïve T cells were labeled with 0.4 µM CFSE in PBS for 10 min at RT. Then the cells were washed with PBS for 3 times. The cells were stimulated with anti-CD3 and anti-CD28 (2 µg/ml) for 48 hours or 72 hours. The cells were collected and stained with anti-CD8. Then the CFSE fluorescence was detected by flow cytometry.

### Measurement of the cytotoxicity of CTL

To measure the cytotoxicity of CTLs, EL-4 cells were pulsed with 10 nM OVA257-264 for 30 min at 37°C. Then the antigen-pulsed EL-4 cells were washed with PBS and then labeled with 1 µM CellTracker Deep Red (CTDR) in serum-free medium for 15 min at 37°C in dark. Meanwhile, EL-4 cells labeled with 0.5 µM CFSE in PBS for 10 min at RT in dark. After washing EL-4 cells with PBS for 3 times, CTDR labeled and CFSE labeled EL-4 cells were mixed at the ratio of 1:1 in the killing medium (RPMI 1640, 2% FBS). CTLs were added into the plate at the ratios of 0:1, 0.5:1, 1:1, 2:1 and 5:1, respectively. After 4 hours, the cytotoxic efficiency was measured by quantifying the value of one minus the ratio of CTDR/CFSE ratio in cytotoxic group to non-cytotoxic group.

### Measurement of the immune synapse formation of CTL

To measure the immune synapse formation between CTL and EL-4 cells, EL-4 cells were pulsed with 10 nM OVA257-264 and labeled with CTDR. CTLs were labeled with CFSE. EL-4 cells and CTLs were mixed at the ratio of 1:1 and co-cultured for 30 min at 37°C. The cells were harvested for flow cytometric analysis and the percentage of CTDR and CFSE double positive cells were quantified.

### LDLR overexpression in CTL

LDLR CDS or D225N mutant sequences were constructed into pMxs-EGFP plasmid. Retrovirus was generated by transfecting platE cells with pMxs-EGFP, pMxs-LDLR-EGFP or pMxs-LDLR D225N-EGFP plasmids. The supernatant containing the retrovirus was collected. To overexpress LDLR in CTL, OT-1 CTLs were generated and cultured for 1 day. Then the cells were spin-infected with the retrovirus for 2 hours at 2000 rpm with 10 ng/ml IL-2 and 10 µg/ml polybrene. Spin-infection was repeated at day 2. EGFP positive cells were isolated by Fluorescence-activated Cell Sorting and cultured in RPMI 1640 complete medium in the presence of 10 ng/ml IL-2.

### Mouse models for colorectal cancer and melanoma

MC38, MC38-OVA or B16F10 cells were washed with PBS and filtered through a 40 µm strainer. Before tumor cells were inoculated, age and sex matched mice (6-8 weeks) were narcotized and shaved first, then 1 × 10^6^ MC38, MC38-OVA cells or 4 × 10^5^ B16F10 cells were subcutaneously injected into the dorsal part of mice. From day 6-10, tumors size was measured every 2 days, and animal survival rate was recorded every day. Tumor size was calculated as length × width. Mice will be euthanized when the tumor size larger than 225mm^2^ (15mm*15mm) for ethical consideration.

### Adoptive T cell transfer

MC38-OVA cells (1 × 10^6^) were injected subcutaneously into *Rag2*^*–/–*^ mice at age 6-8 weeks. On day 12-14, tumor-bearing mice with similar tumor size were randomly divided into specific groups and respectively received PBS, wild-type OT-I CTLs (1 × 10^6^), *Ldlr*^*–/–*^ OT-I CTLs (1 × 10^6^) or *PCSK9*^*–/–*^ OT-I CTLs (1 × 10^6^) intravenously injection. Tumor size was calculated as length × width every 2 days and animal survival was measured every day from day 8. When the tumor size was larger than 225 mm^2^, the mice were euthanized for ethical consideration.

### Depletion of CD8^+^ T cells

Mc38 cells (1 × 10^6^) were inoculated subcutaneously into C57BL/6 mice at 6-8 weeks. Two days before tumor inoculation, 200 µg/ml of α-CD8 depletion antibody (2.43, Bio X Cell) or rat IgG (2A3, Bio X Cell) were intraperitoneally injected into indicated group. Subsequently, α-CD8 depletion antibody or rat IgG were injected for every 4 days.

### Treatment of cancer with PF-06446846, anti-PD-1 antibody or PF-06446846 plus anti-PD-1 antibody in vivo

Tumor-bearing mice with similar tumor size were randomly divided into different groups and received PBS, anti-PD-1 antibody (RMP1-14, Bio X Cell, 200 µg per injection), PF-06446846 (2 mg/ml, 5 mg/ml or 10 mg/ml as indicated) or anti-PD-1 antibody plus PF-06446846 injection intraperitoneally every 2 days, respectively. PF-06446846 was injected 8 times from day 8 and anti-PD-1 was injected 6 times from day 9. The tumor size and survival were measured as mentioned above.

### Tumor infiltrating lymphocytes isolation and analysis

CTL adoptively transferred *Rag2*^-/-^ mice or tumor-bearing C57BL/6 mice were anesthetized and sacrificed, tumor tissues were dissected and cut into pieces and digested in RPMI 1640 medium containing collagenase VI (210 U/ml), DNase I (100 U/ml) and hyaluronidase (0.5 mg/ml) for 30 min at 37°C. The dissociated cells were passed through a 70 µm strainer. The filtered cells were centrifuged at 50 g for 1 min. Then the supernatant was removed to a new tube to centrifuge at 1000 g for 10 min. Resuspended cells for density gradient centrifugation with 40% Percoll and 70% Percoll. Harvested the interphase of gradient and spin at 1000 g for 5 min. The isolated tumor infiltrated lymphocytes were then to do the subsequent experiments. To measure the cytokine production of isolated TILs, the cells were stimulated with 50 ng/ml PMA, 1 µM ionomycin and 5 µg/ml BFA for 4 hours at 37 °C.

### CD8^+^ T cells selection

Isolate CD8^+^ T cells in tumor infiltrated lymphocytes was based on EasySep™ Release Mouse Biotin Positive Selection Kit (Stemcell). In brief, tumor infiltrated lymphocytes were resuspended in 500 µl (5 × 10^7), added biotin labeled anti-mouse CD8 (53-6.7) antibody and incubated for 15 min at RT. Washed cells with isolation buffer and centrifuge for 5 min at 400 g. Added selection cocktail and incubated for 15 min at RT. Then RapidSpheres beads were added into incubation system for 10 min at RT under rolling and tilting. After incubating, add isolation buffer and magnetically select bead-bound CD8^+^ T cells. Washed bead-bounded CD8^+^ T cells for 3 times and obtain pure bead-bounded CD8^+^ T cells.

### Filipin III staining

Isolated tumor infiltrated T cells were washed with PBS for 3 times. Then load cells on the glass dish and incubate at RT for 10 min. Add 4% paraformaldehyde (PFA) and 0.05% glutaraldehyde to fix cells at RT for 10 minutes. Wash cells with PBS for 3 times and then stain Filipin III at the concentration of 50 ng/ml for 2 hours at RT. Cells were washed for 8 times and images were collected using Zeiss (LSM880, AxioObserver) confocal microscope and analyzed using Image J software.

### Modulation of the plasma membrane cholesterol level by MβCD and MβCD-coated cholesterol

To deplete cholesterol from the plasma membrane, CD8^+^ T cells were washed with PBS for two times and then incubated with 1 mM MβCD at 37 °C for 15 min. The cells were then washed three times with PBS.

To add cholesterol to the plasma membrane, CD8^+^ T cells were washed with PBS for two times and then incubated with 10 μg/ml MβCD-coated cholesterol at 37 °C for 15 min. The cells were then washed three times with PBS.

### PCSK9 and PF-06446846 treatment

Isolated naïve CD8^+^ T cells from the spleen were stimulated with anti-CD3 and anti-CD28 in the presence of PCSK9 (5 μg/ml) for 24 hours and cytokine production were then determined.

EL4 and EL4-OVA cells were pretreated with PF-06446846 (5 μM or 10 μM) for 24 hours and then cocultured with CTLs for 12 hours. The cytotoxic efficiency was measured by flow cytometry.

### Immunofluoresence detection of co-localization and immune synapse

CTLs were harvested and placed in glass bottom cell culture dish and fixed with 4% PFA. After blocking the non-specific binding sites with goat serum for 30 min at RT, the cells were incubated with anti-LDLR (Lifespan) and anti-CD3 (Genetex) primary antibodies for 12 hours at 4°C. Then the cells were stained with Alexa 488-conjugated goat anti-rabbit IgG and Alexa Fluor Plus 555-conjugated donkey anti-mouse IgG for 2 hours at 4°C after washing with PBS. Before imaging, the cells were sealed with In Situ Mounting Medium with DAPI (Sigma). Images were collected using Zessi (LSM880, AxioObserver) confocal microscope.

For immune synapse detection, LDLR-mEOS3.2 overexpressed CTLs and MC38-OVA were resuspended 4:1 with serum-free RPMI-1640 at 1 million per milliliter. Cells were centrifuged at 200 rpm for 1 min, then incubated at 37°C for 15 min. Then cells were plated in glass bottom cell culture dish and allowed to settle down for 15 min before fixation with 4% PFA for 10 min. The cells were blocked with goat serum for 60 min at RT, stained with anti-CD3ε (Bio X Cell, 1ug/ml) primary antibody for 12 hours at 4°C. Excessive antibody was washed with PBS for three times. Then the cells were stained with Alexa 647-conjugated goat anti-hamster IgG secondary antibody for 60 min at RT. After washing with PBS, the cells were sealed with DAPI containing anti-fade reagent (Sigma). Images were collected with Nikon N-SIM system.

### Measurement the interaction of LDLR and CD3 by proximity ligation assay (PLA)

PLA allows for endogenous detection of protein interaction. We detect the interaction of LDLR and CD3 according to Duolink PLA Fluorescence protocol (Sigma). Jurkat cells, activated Jurkat cells, CD8^+^ T cells from wild type mice and *Ldlr*^*–/–*^ mice were loaded on glass dish. Cells were fixed with 4% PFA. Block non-specific signal by adding Duolink Blocking Solution and incubate for 60 min at 37°C. After blocking, add the anti-LDLR and anti-CD3 primary antibodies and incubated for 12h at 4°C. Then two PLA probes were diluted and added to the samples and incubated for 60 min at 37°C. Prepare ligation and amplification buffer to ligate the fluorescence probe and amplify the signal. Mount the samples with In Situ Mounting Medium with DAPI (Sigma). The images were captured with Olympus FV1000 or Zess LSM880 confocal microscope, and analyzed with Image J software.

The TIRF-imaging was performed on Nikon N-SIM + N-STORM microscope with an TIRF 100 × oil immersion lens. Adjusted the oblique incidence excitation to the appropriate TIRF angle to capture images.

### Co-immunoprecipitation and western blot analysis

EL4 cells and CTLs were lysed in Nonidet P-40 lysis buffer (50 mM Tris-HCl, pH 7.4, 155 mM NaCl, 5 mM EDTA, 2 mM Na3VO4, 20 mM NaF, supplemented with complete protease inhibitor cocktail and phosphatase inhibitor cocktail), and target protein was immunoprecipitated with corresponding antibody and by Pierce™ Co-Immunoprecipitation Kit (Thermo Fisher) according to the manufacturer’s instructions.

For western blotting analysis, proteins were separated by SDS-PAGE and transferred to polyvinyldifluoride (PVDF) membrane. Proteins were then probed with specific primary antibodies followed by secondary antibodies conjugated with horseradish percidase (HRP).

### Statistics

Statistical parameters are all shown in Figure Legends. Statistical analysis was performed using nonparametric two-tailed *t* test or two-way ANOVA in GraphPad Prism. The survival were analyzed by using Log-rank (Mantel-Cox) test. Unless specially described, error bars stand for standard error of the mean. *, *P* < 0.05; **, *P* < 0.01; ***, *P* < 0.001; ****, *P* < 0.0001.

## Acknowledgements

The imaging works were performed at the SMU Central Laboratory of Southern Medical University and Department of Pathology of Nanfang Hospital. We thank Chenqi Xu for the discussion at the early stage of this project. W.Y. is funded by National Key R&D Program of China (MOST, No. 2018YFA0800404), NSFC grants (No. 81822036 and 31770931), Guangdong Natural Science Funds for Distinguished Young Scholar (No. 2017A030306030). J.Y. is funded by NSFC grants (No. 82001658), China Postdoctoral Science Foundation (No. BX20190148 and 2019M662973) and Guangdaong Basic and Applied Basic Research Foundation (No. 2019A1515110015). T.C. is funded by NSFC grants (No. 82001745), China Postdoctoral Science Foundation (No. 2020M672544) and Guangdaong Basic and Applied Basic Research Foundation (No. 2019A1515110052). X.Z. is funded by NSFC grants (No. 31800730), China Postdoctoral Science Foundation (No. 2017M622730), Natural Science Foundation of Guangdong Province (No. 2018030310293) and Guangdaong Basic and Applied Basic Research Foundation (No. 2020A1515011246).

## Author contributions

W.Y. conceived the project. W.Y., Y.D., H.Z., J.Y., T.C. and X.Z. designed the experiments; J.Y., X.Z., H.L. and Q.C. performed the cellular experiments; T.C., J.Y., Z.C., H.C. and M.L. performed the animal experiments; X.Z., J.Y. and X.L. performed the biochemical and imaging experiment; Y.R., J.Q., S.F., and Y.W performed the IHC experiments and analyzed the clinical samples; Y.D. and H.Z. contributed to direct the project and discussions. All the authors contributed to data analysis, manuscript writing and revision.

---

**Supplementary figure 1.**
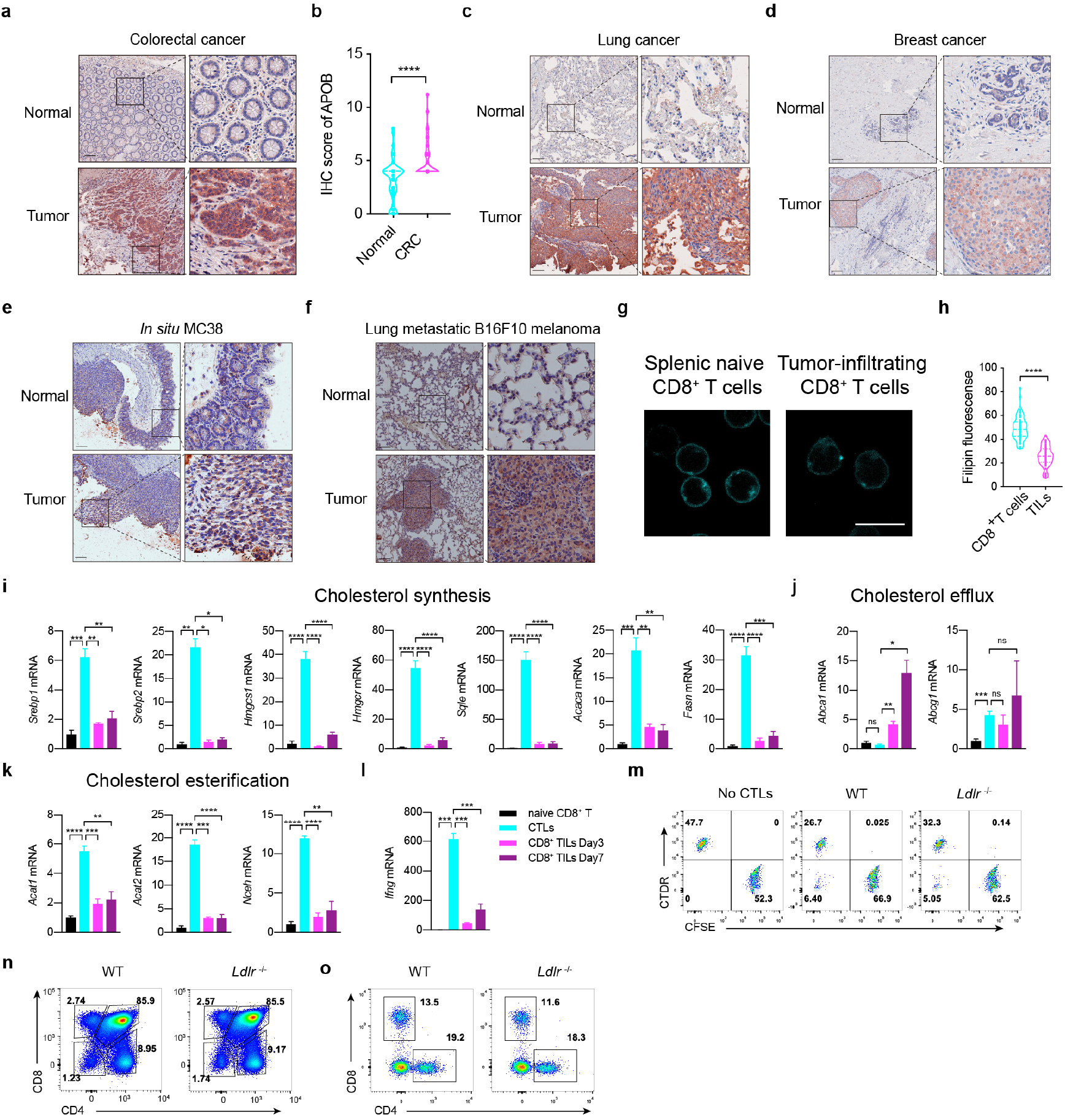
LDLR deficiency hinders the antitumor activity of CD8^+^ T cells. (related to Figure 1) **a-b**, Human normal colorectal sections or tumor sections were stained with anti-APOB antibody by immunohistochemistry and the abundance of APOB was assessed in (**b**), (n = 50). **c**, Human normal lung or tumor sections were stained with anti-APOB antibody by immunohistochemistry. **d**, Human normal breast or tumor sections were stained with anti-APOB antibody by immunohistochemistry. **e**, MC38 cells were injected into the cecum of C57BL/6 mice and the tumor sections were stained with anti-APOB antibody by immunohistochemistry. **f**, B16F10 cells were intravenously injected into C57BL/6 mice to induce the lung metastasis of melanoma. The tumor sections were stained with anti-APOB antibody by immunohistochemistry. Scale bar, 120μm(**a-f**). **g-h**, Filipin III staining to analyse cellular cholesterol distribution in splenic naive and tumor-infiltrating CD8^+^ T cells. The Filipin fluorescence was analyzed in (h). Scale bar, 10μm. **i-l**, Transcriptional levels of cholesterol synthesis (**i**), efflux (**j**), esterification (**k**) and *Ifng* as a control were analyzed by QPCR in naïve CD8 T cells, CTL and CD8^+^ TILs (isolated at Day3 or Day7 post adoptive transfer), (n = 4). **m**, Cytotoxicity of WT and *Ldlr*^-/-^ CTLs. WT and *Ldlr*^-/-^ OT-I CTLs were incubated with OVA-pulsed CTDR-labeled EL-4 cells and CFSE-labeled non-pulsed EL-4 cells for 4 hours. **n-o**, T cell development analysis of thymocytes (**n**) and splenic (**o**) T cells of WT and *Ldlr*^-/-^ mice. *, *P* < 0.05; **, *P* < 0.01; ***, *P* < 0.001****, *P* < 0.0001. Error bars denote for the s.e.m.

**Supplementary figure 2.**
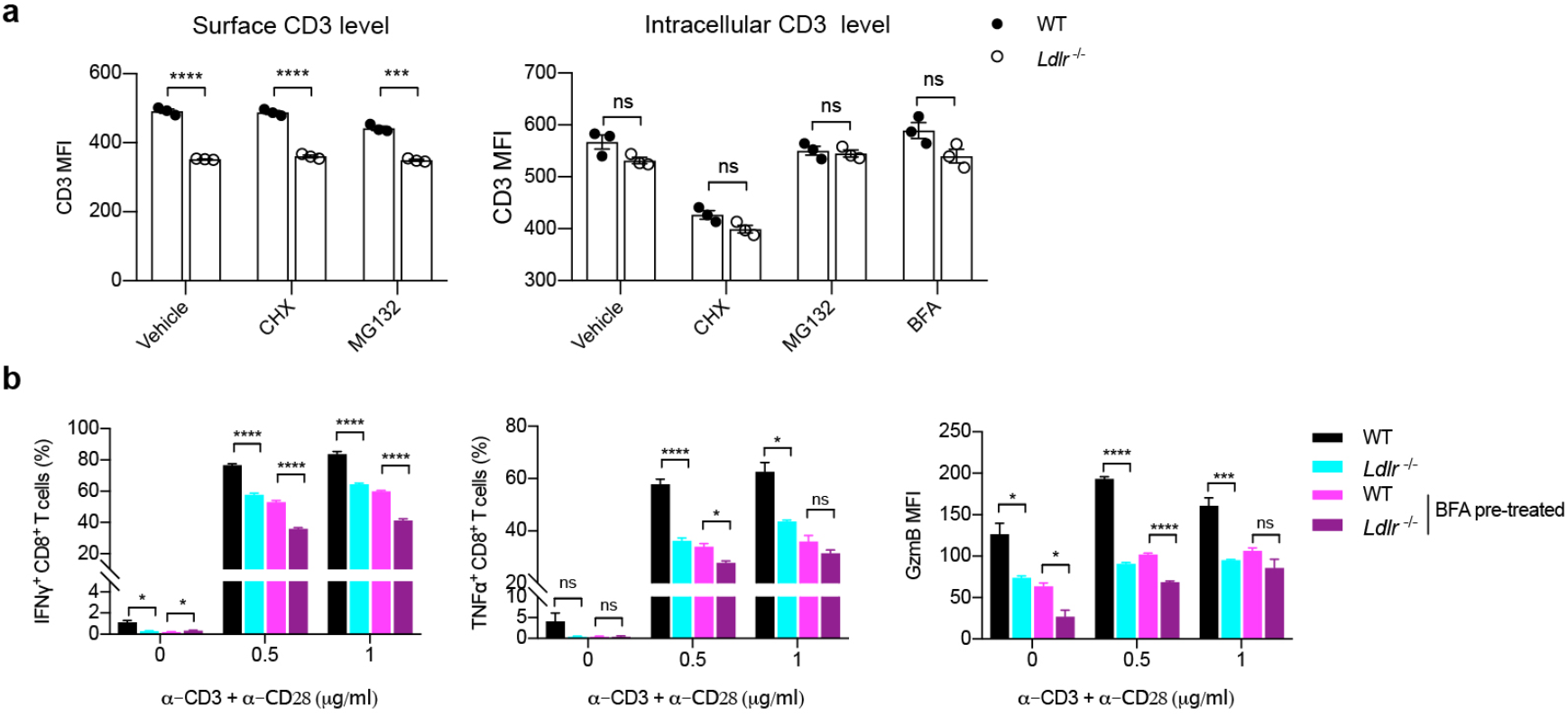
LDLR binds to TCR and regulate TCR signaling in CD8^+^ T cells. (related to Figure 3) **a**, CTLs were treated with CHX (50μg/ml), MG132 (15μM), BFA (5μg/ml) or not for 2 hours. Surface levels (left panel) and intracellular levels (right panel) of CD3 were analyzed by flow cytometry. Data were analyzed by *t* test (n = 3). **b**, Cytokine and granule productions of WT and *Ldlr*^-/-^ CTLs. CTLs were pretreated with BFA (5μg/ml) or not for 2 hours and stimulated with anti-CD3 and anti-CD28 antibodies for 4 hours at indicated concentrations. Data were analyzed by *t* test (n = 4). ns, no significance; *, *P* < 0.05; ***, *P* < 0.001; ****, *P* <0.0001. Error bars denote for the s.e.m.

**Supplementary figure 3.**
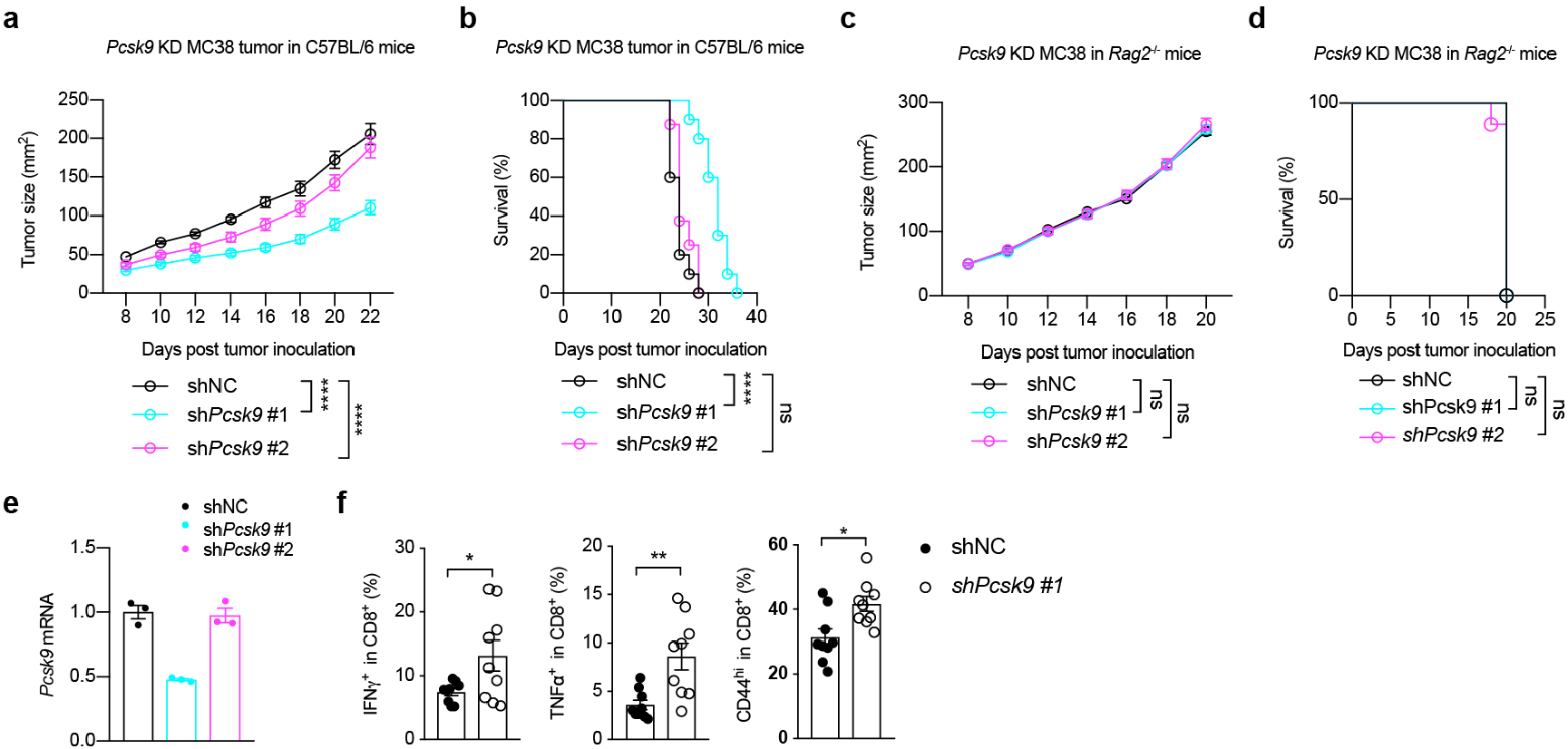
Tumor-derived PCSK9 inhibits the antitumor activity of CD8^+^ T cells (related to Figure 4) **a, b**, Tumor growth (**a**) and survival (**b**) of *Pcsk9* knockdown MC38 tumor-bearing C57BL/6 mice. Data were analyzed by two-way ANOVA (n = 8-10). **c, d**, Tumor growth (**c**) and survival (**d**) of *Pcsk9* knockdown MC38 tumor-bearing Rag2^-/-^ mice. Data were analyzed by two-way ANOVA (n = 9). **e**, Transcriptional level of *Pcsk9* was measured by QPCR in *Pcsk9* knockdown MC38 cells.**f**, Cytokine productions and activation of tumor infiltration CD8^+^ T cells isolated from shNC or sh*Pcsk9* MC38 tumors. Data were analyzed by *t* test (n = 9). ns, no significance; *, *P* < 0.05; **, *P* < 0.01; ****, *P* <0.0001. Error bars denote for the s.e.m.

**Supplementary figure 4.**
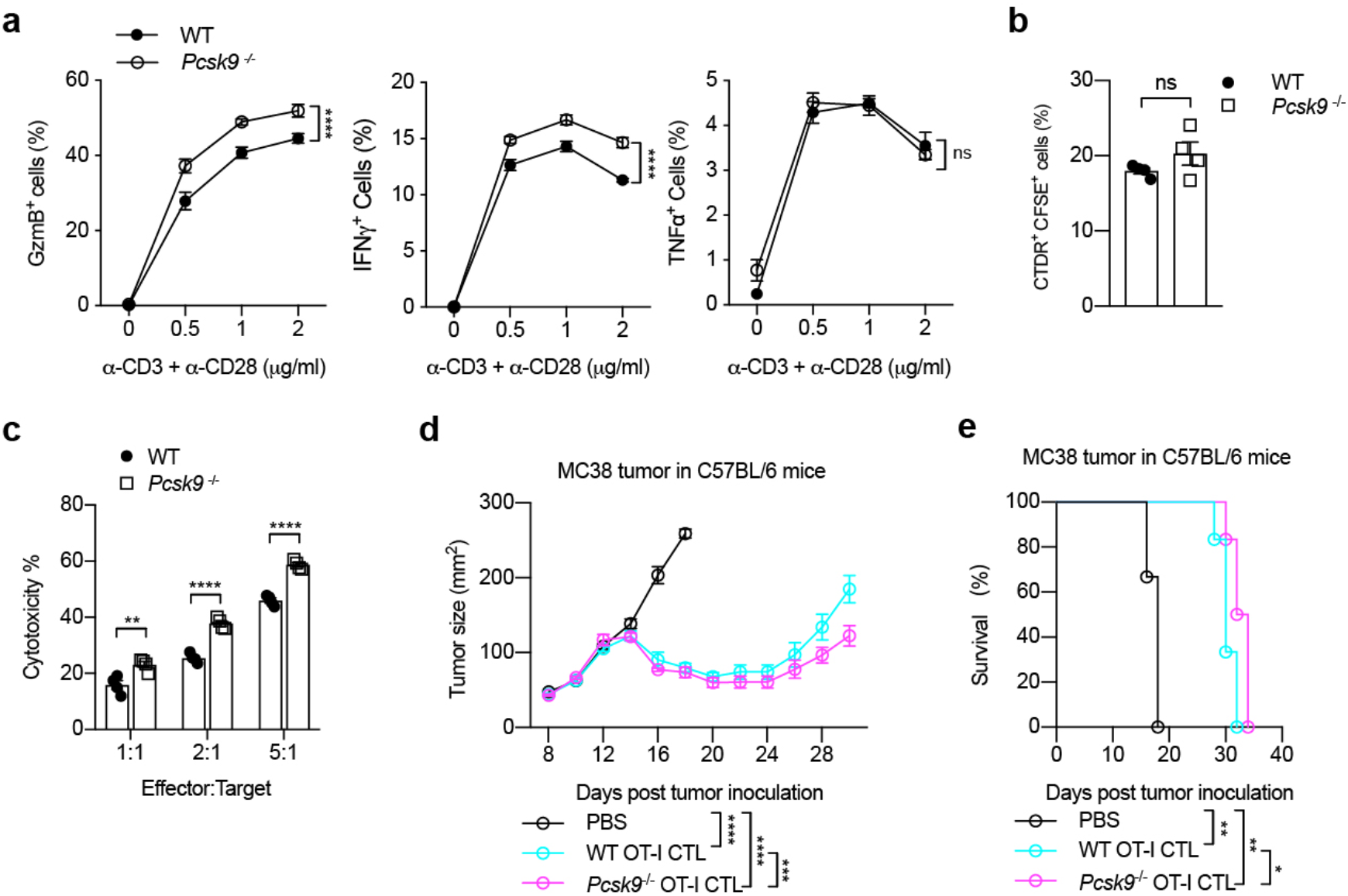
Inhibiting PCSK9 potentiating the antitumor activity of CD8+ T cells (related to Figure 5) **a**, Cytokine/granule productions of WT and *Pcsk9*^-/-^ CD8^+^ T cells. CD8^+^ T cells were isolated from the spleen of WT or *Pcsk9*^-/-^ mice and stimulated with anti-CD3 and anti-CD28 antibodies at indicated concentrations for 24 hours. **b**, Synapse formation of WT and *Pcsk9*^*-/-*^ CTLs. CFSE-labeled CTLs and CTDR-labeled OVA-pulsed EL4 cells were cocultured for 30 mins. Data were analyzed by *t* test (n = 4). **c**, Cytotoxicity of WT and *Pcsk9*^*-/-*^ CTLs. CTLs were incubated with OVA-pulsed CTDR-labeled EL-4 cells and CFSE-labeled non-pulsed EL-4 cells for 4 hours. Data were analyzed by *t* test (n = 4). **d, e**, Tumor growth (d) and survival (e) of MC38-OVA tumor-bearing Rag2^-/-^ mice after adoptive transfer of PBS, WT or *Pcsk9*^*-/-*^ CTLs. Data were analyzed by two-way ANOVA (n = 6). ns, no significance; *, *P* < 0.05; **, *P* < 0.01; ***, *P* < 0.001; ****, *P* <0.0001. Error bars denote for the s.e.m.

**Supplementary figure 5.**
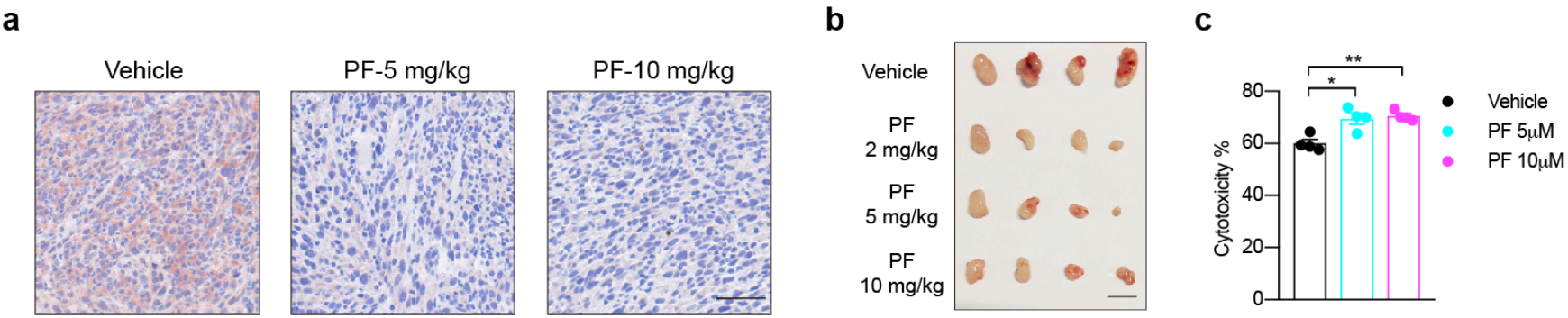
Inhibiting PCSK9 potentiating the antitumor activity of CD8+ T cells (related to Figure 6) **a**, PCSK9 expression was measured in MC38 tumor sections by immunohistochemistry. MC38 cells were subcutaneously injected into C57BL/6 mice and treated intraperitoneally with PCSK9 inhibitor PF0644684 every 2 days at indicated concentrations. Scale bar, 50μm. **b**, MC38 tumors were isolated from PF0644684 treated C57BL/6 mice and tumor size was shown. Scale bar, 10mm.**c**, Cytotoxicity of CTLs cocultured with PF0644684 treated EL4 cells. EL4 cells were pretreated with PF0644684 for 24 hours and cocultured with CTLs for 12 hours in the presence of PF0644684. Data were analyzed by *t* test (n = 4). *, *P* < 0.05; **, *P* < 0.01. Error bars denote for the s.e.m.

